# On-Off coding is latent in vertebrate visual circuits

**DOI:** 10.64898/2026.05.01.722149

**Authors:** Xinwei Wang, Laura Steel, Chiara Fornetto, Dominic Gonschorek, Marvin Seifert, Timm Schubert, Stephan CF Neuhauss, Thomas Euler, Tom Baden

**Author notes:** Correspondence: XW; TB. equal second authors (Laura Steel, Chiara Fornetto, Dominic Gonschorek).

## Abstract

Across sensory systems, neurons often encode stimulus changes of opposite sign as On and Off signals, a fundamental strategy for representing deviations from baseline. In vertebrate vision, this computation is classically attributed to an early, hardwired split in the retina, with segregated pathways subsequently propagated through downstream circuits. Here, we challenge this view. Combining comparative transcriptomics, pharmacology, genetics, in vivo imaging and electrophysiology in zebrafish, with validation in mouse retina, we show that On-Off coding is a latent and intrinsic property of visual circuits. Bipolar cells frequently co-express receptor systems of opposing polarity and can generate mixed responses that are normally suppressed by inhibitory circuitry. Disrupting inhibition or selectively blocking receptor pathways unmasks robust On-Off signalling. In addition, mixed On-Off responses emerge naturally as effective light input increases, including during development and at higher light levels. Across processing stages, including retinal ganglion cells and central neurons, inhibition repeatedly enforces apparent polarity segregation. Thus, polarity splitting is a distributed, dynamically regulated computation rather than a fixed circuit feature.

**HIGHLIGHTS:**

- Mixed On-Off signals are latent across visual processing stages
- Inhibition actively sculpts polarity, not just refines it
- Polarity splitting is a dynamic, distributed and reusable computation
- Conserved mechanism revealed in zebrafish and mouse

## INTRODUCTION

Across sensory systems, neural circuits routinely represent stimulus changes of opposite sign, classically termed On and Off signals^1^. Across both vertebrate and lineages, such polarity-split representations are found in vision^2–4^, olfaction^5,6^, mechanosensation^7,8^, and audition^9,10^. Their repeated emergence suggests deep computational value, including expanded dynamic range^11^, efficient encoding of deviations from baseline^12,13^, decorrelation^14,15^, and as a stepping stone for downstream computations^1,16^.

A dominant intuition is that polarity splitting is achieved through a specific, early circuit transformation, after which On and Off signals are largely propagated rather than reconstructed. In vertebrate vision, this view is grounded in the canonical mammalian retinal circuit^16^. Photoreceptors release glutamate^17^ tonically in darkness and hyperpolarise to light, rendering them functionally Off^18^. One synapse downstream, retinal bipolar cells (BCs) segregate into On and Off types via a molecular dichotomy^19^: Off BCs express sign-preserving ionotropic AMPA/kainate receptors^20–22^, whereas On BCs express the sign-inverting metabotropic receptor mGluR6^19^, which closes TRPM1 channels through a G-protein cascade^23^. This scheme has become the textbook model of retinal pathway splitting^24^ and has strongly influenced thinking across sensory neuroscience.

Implicit in this framework is the idea that most downstream circuits inherit segregated polarity channels^16^. Crossover inhibition^25^, the spread of opposite polarity inhibitory signals between Off and On circuits, is thought to further stabilise this ‘pre-existing’ condition. Consistent with this, many mammalian retinal ganglion cells (RGCs) are classified as predominantly On or Off^26–31^, and mixed responses are often treated as special-purpose exceptions^32,33^ that arise from the convergence of otherwise separate pathways^34^.

However, this segregation is not absolute. For example, in mice, On-Off responses become more prominent at higher light levels^35^ and can be dynamically accentuated or suppressed under a wide range of experimental manipulations^14,36–45^. Bipolar cells also do not always conform strictly to the dichotomy^14,46–48^: For example, anatomical stratification can blur canonical boundaries^49^. Comparative evidence further suggests ecological modulation, with secondarily diurnal mammals exhibiting stronger On-Off coding than nocturnal or crepuscular rodents^50,51^.

The view of largely ‘fixed’ On versus Off circuits is even harder to reconcile with non-mammalian vertebrates, including birds^52^, reptiles^53^, amphibians^54–56^, and teleost fish^57^, where the retinal output is routinely dominated by RGCs with mixed On-Off responses. Rather than a special feature of select motifs, On-Off coding appears to be a prevalent mode of retinal output in these species. Moreover, bona fide On-Off bipolar cells are present in several of these species^39,58–62^, and polarity inversion is already evident at the photoreceptors via horizontal-cell–driven network interactions^63,64^, indicating that mixed polarity can emerge at the earliest synaptic stages. Such signals are not merely tolerated but actively exploited to multiplex visual features within single output channels^52,65^, emphasising On-Off coding as a rich representational substrate.

These observations motivate an alternative hypothesis: that On-Off coding is not primarily created by merging segregated pathways but is instead a latent and broadly available property of visual circuits. Here, we use ‘latent’ to denote signals that are present in the circuit but not expressed due to active suppression. Under this view, neurons are fundamentally capable of representing both polarities, with apparent unipolarity emerging when one sign is selectively suppressed. Polarity segregation would therefore not be solved once at the first synapse but repeatedly reconstructed and actively maintained at successive processing stages, allowing circuits to flexibly deploy either mixed or segregated codes depending on computational demands.

A key implication of this framework is that polarity coding should not be static, but dynamically adjustable. In particular, the balance between mixed and segregated representations may depend on the effective signal-to-noise regime, which varies with light level, contrast, and photon catch. Under high-signal conditions, circuits may increasingly exploit mixed On-Off representations, whereas under low-signal conditions, stronger polarity segregation may stabilise responses. Such a regime-dependent shift has been suggested in mammalian retina^35^, but whether it reflects a general and intrinsic property of visual circuits remains unclear.

Here, we test this hypothesis by examining polarity coding in the zebrafish visual system, a primitively diurnal, cone-dominated vertebrate model that preserves many ancestral visual features^66–68^. Using comparative transcriptomics, pharmacology, genetics, *in vivo* functional imaging, and multielectrode array recordings, we show that On-Off coding is fundamental, widespread, and repeatedly reconfigured across processing stages. Many bipolar cells are intrinsically On-Off, co-expressing receptors of opposing polarity, and GABAergic inhibitory circuits suppress one sign to enforce apparent segregation. Strikingly, the same logic recurs downstream: blocking inhibition unmasks strong On-Off responses in RGCs and again in central visual circuits. GABAergic inhibition block also unmasks latent On-Off responses in mouse RGCs. Together, our results reveal On-Off coding as a latent representational mode that is actively sculpted and flexibly redeployed at multiple stages of the visual pathway, with similar downstream mechanisms present across vertebrates.

## RESULTS

### On-Off polarity is latent at the level of retinal bipolar cells

The prevailing model of retinal pathway splitting assumes that bipolar cells (BCs) are intrinsically unipolar, either On or Off, by virtue of their glutamate receptor complement at the photoreceptor synapse^16^. However, several studies in non-mammalian retinas have reported BCs with robust mixed On-Off responses^39,58–62^, and transcriptomic studies across vertebrates reveal extensive co-expression of receptors associated with opposite polarities^69–71^. Together, these observations raise the possibility that bipolar cell mixed-polarity signalling may be widespread but normally masked by circuit mechanisms.

Across species, we therefore assessed how consistently receptor identity alone predicts ‘transcriptomic polarity’, which is defined by deeply conserved transcriptional programs and broader cell-type expression signatures rather than receptor choice alone^70,71^. To this end, we surveyed available single cell RNA sequencing (snRNAseq) datasets for glutamate receptor expression in retinal BCs across vertebrates^69–71^. We focussed on ionotropic kainate and AMPA receptors as sign-preserving (Off) mechanisms, and mGluR6 and excitatory amino acid transporters (EAATs)^61,72,73^ as sign-inverting (On) mechanisms^16^ (Fig. 1A). Across species, two general features emerged. First, receptor expression only weakly predicts ‘transcriptomic polarity’. Second, systematic differences in receptor usage correlate with phylogeny and ecology, particularly mammalian identity and rod dominance.

**Figure 1.**
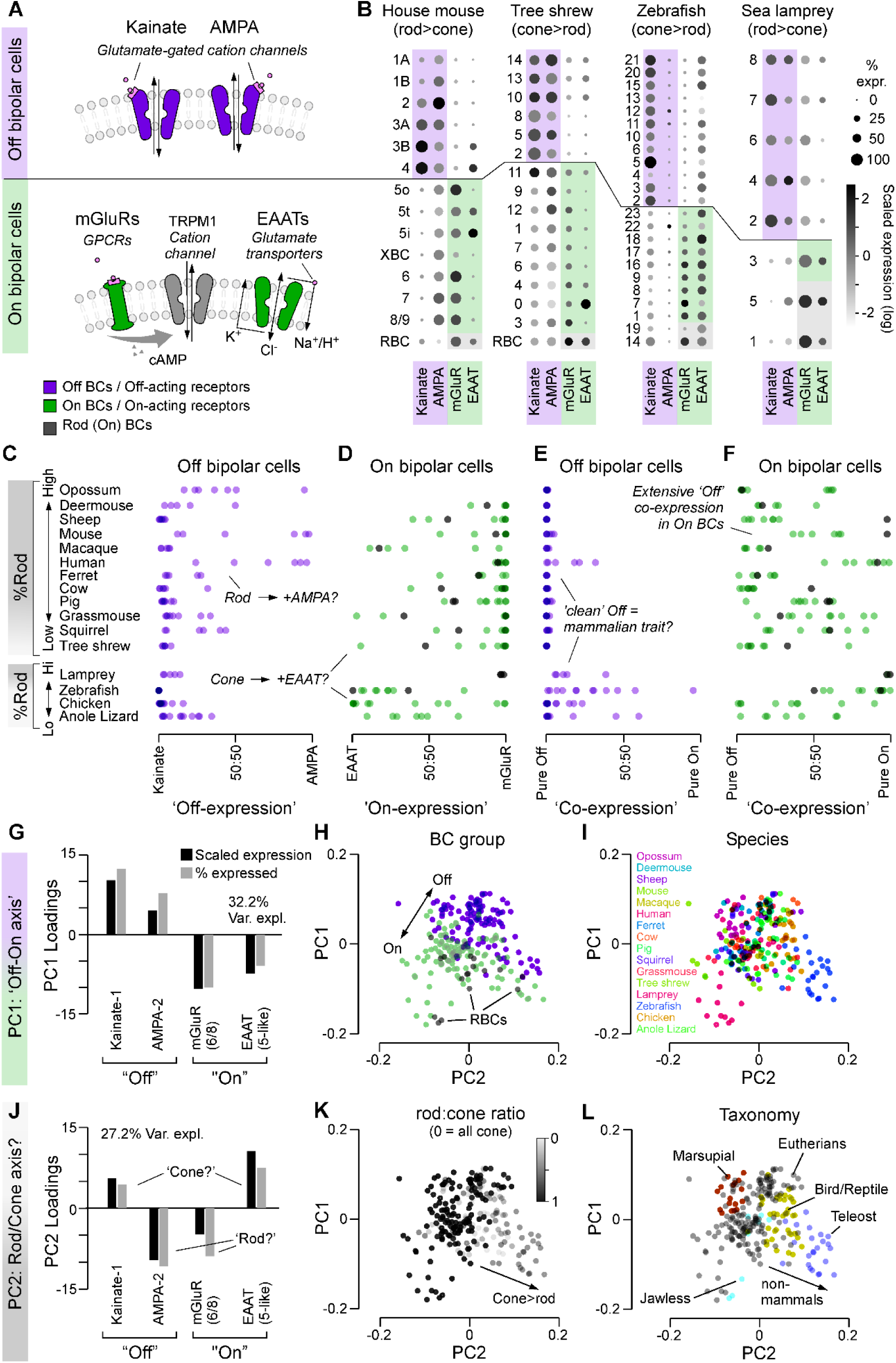
Comparative transcriptomics of outer retinal glutamate receptors. **A,B**, Major classes of Off-acting (kainate and AMPA) and On-acting (mGluRs and excitatory amino acid transporters, EAATs) glutamate receptors in retinal bipolar cells (BCs) (**A**), and their scRNA-seq-based expression profiles in transcriptomically defined On and Off BC types from mouse, tree shrew, zebrafish, and sea lamprey (**B**; data from Refs^69–71,110^). In general, “mGluR” refers to grm6, whereas “EAAT” refers to EAAT5-like genes such as slc1a7. Because gene names, orthologues, and expression patterns are not perfectly aligned across species, the terms kainate, AMPA, mGluR, and EAAT refer here to the most relevant gene sets in each species (see Supplemental Table T1 for correspondence). Additional species are shown in Supplemental Figure S1A. **C–F,** Expression contrasts quantifying the relative expression levels indicated for each BC cluster (Methods): kainate versus AMPA in Off BCs (**C**), EAAT versus mGluR in On BCs (**D**), Σ(kainate, AMPA) versus Σ(mGluR, EAAT) in Off BCs (**E**), and Σ(kainate, AMPA) versus Σ(mGluR, EAAT) in On BCs (**F**). Rod bipolar cell orthotypes^69,70^ are colour-coded in grey and included within the On-BC group. Species are split by taxonomy (mammals versus non-mammals) and ordered by rod dominance (cf. Supplemental Figure S1B). **G–L,** Principal component analysis (PCA) of BC glutamate receptor expression patterns across species (Methods), showing loadings of the first (**G**) and second (**J**) principal components, and the distribution of BCs in PCA space colour-coded by BC group (**H**), species (**I**), rod:cone ratio (**K**), and taxonomy (**L**; see also Supplemental Figure S1C).

In the rod-dominated house mouse, kainate receptors and mGluR6 align relatively closely with Off and On BCs, respectively (Fig. 1B). However, AMPA receptors and EAATs appear indiscriminately present across both polarity groups. Such mixed receptor presence becomes even more pronounced in some secondarily diurnal mammals such as tree shrew, grass mouse or pig (cf. Supplemental Figure 1A). In the former, for example, at least one On BC type robustly expresses kainate receptors, and mGluR6 expression is weaker and less selectively confined to On types. Comparable intermixing is observed in non-mammalian vertebrates. For example, in zebrafish, AMPA receptor expression is low and EAAT5b expression is high across most BC types.

To systematically quantify these trends, we analysed available datasets^70^ from twelve mammals and four non-mammalian vertebrates (lamprey, zebrafish, chicken, anole lizard), ranked by rod-to-cone ratio (Supplementary Fig. S1B). For each receptor gene (Supplementary Table T1), we computed an expression factor incorporating both prevalence and expression level and quantified the relative contributions of kainate versus AMPA receptors in Off BCs, and EAATs versus mGluR6 in On BCs (Fig. 1C-F). This analysis revealed several patterns. First, non-mammalian species were predominantly kainate-dominated in the Off pathway, whereas some highly rod-dominated mammals additionally employed AMPA receptors (Fig. 1C).

Conversely, mGluR6 dominance was most pronounced in mammals and correlated with rod-dominance, while EAAT-mediated On signalling was enriched in cone-dominated species (Fig. 1D). In some species this segregation appears to be more strongly biased at the level of receptor expression, with On- and Off-bipolar cell populations showing relatively limited overlap in their associated receptor repertoires. This suggests that, in these cases, polarity separation is at least partly supported by molecular specialisation.

We next quantified receptor co-expression by measuring the presence of On-associated receptors in Off BCs and Off-associated receptors in On BCs (Fig. 1E,F). Most species contained substantial populations of BCs that robustly co-expressed receptors of opposing polarity. Mammalian Off BCs were generally ‘clean’, whereas non-mammalian Off BCs frequently co-expressed On-associated receptors (Fig. 1E). By contrast, On BCs showed extensive heterogeneity across all species, with widespread co-expression of Off-associated receptors (Fig. 1F).

An unbiased principal component analysis captured these relationships (Fig. 1G-L): the first component separated receptors by polarity (On versus Off, Fig. 1G,H), largely independent of species (Fig. 1I), while the second contrasted distinct receptor strategies (kainate + EAAT versus AMPA + mGluR6, Fig. 1J) and aligned with both rod–cone ratio (Fig. 1K) and taxonomic grouping (Fig. 1L).

Together, these analyses indicate that molecular receptor identity alone cannot fully explain transcriptomic polarity. Instead, they suggest that the clean On versus Off segregation classically described in some mammals may represent a derived specialisation rather than a general vertebrate strategy.

To understand how polarity is organised in a diurnal, non-mammalian retina that more closely reflects the inferred ancestral input circuitry^68,74^, we therefore turned to zebrafish^66^. We used 2-photon imaging to assess the polarity of visual responses in the retina and brain of larval zebrafish (6-8 days post fertilisation, *dpf*), in the context of pharmacological and/or genetic manipulations that selectively interfered with distinct circuit elements in the visual pathway. Throughout, we used spatially widefield and spectrally broad steps of light for visual stimulation (Methods). We began with bipolar cells.

### Inner retinal inhibition suppresses intrinsic On-Off bipolar cell signalling

If bipolar cells are capable of mixed-polarity signalling, then removing circuit mechanisms that normally enforce polarity segregation, should permit such responses to be observed^75^. For example, under control conditions, bipolar cell terminals exhibit Off-dominated responses in the ‘upper’ inner plexiform layer (IPL) and On-dominated responses in the ‘lower’ IPL (Fig. 2A,B). However, following pharmacological removal of both GABA- and glycinergic inhibitory input from amacrine cells using GABAzine, TPMPA and strychnine (Methods), approximately one third of terminals exhibited clear responses at both contrast polarities (Fig. 2C-F, cf. Supplemental Movie M1). Most of these latent On-Off terminals were observed within two reproducible IPL strata (Fig. 2G), indicating that they belong to specific bipolar cell types^69,76^. This observation is consistent with our transcriptomic analysis (Fig. 1), which revealed widespread co-expression of receptor systems associated with opposing polarities.

**Figure 2.**
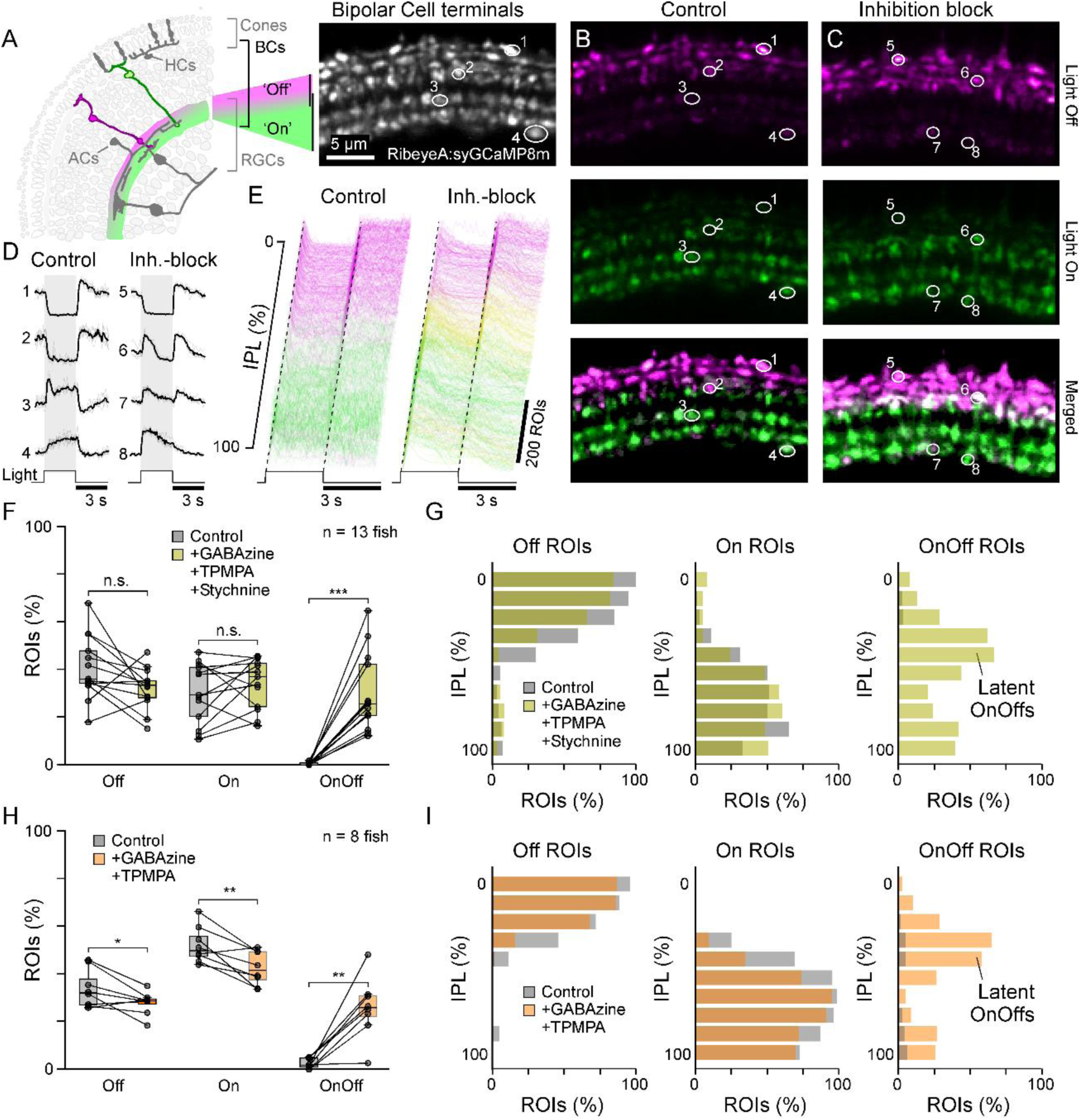
Blocking inner retinal inhibition unmasks latent On-Off bipolar cells. **A–D**, Example 2-photon recordings of retinal bipolar cell terminals in the inner plexiform layer (IPL) of larval zebrafish (6-8 days post fertilisation, dpf) in response to a 3-s “white” light step (Methods). Shown are the recorded region (**A**), the control condition (**B**), and a nearby retinal region from the same fish following intraocular injection of an inhibitory cocktail blocking GABAA (Gabazine), GABAC (TPMPA), and glycine receptors (strychnine) (**C**, inhibition block). **E,** Summary of 750 BC responses, subsampled from n = 1,527 ROIs (control) and n = 1,639 ROIs (inhibition block) across 13 fish, arranged by vertical position in the IPL. For visualisation, responses classified as Off, On-Off, and On (Methods) are colour-coded magenta, yellow, and green, respectively. **F,G,** Summary of the distributions of On, Off, and On-Off responses across fish (**F**; statistics: paired t-test; see Supplemental Table T2) and across IPL depth (**G**). **H,I,** As in **F,G**, but for block of GABAA and GABAC receptors only (n = 1,204 and 1,381 ROIs, control/drug; n = 8 fish). Single-drug manipulations are shown in Supplemental Figure S2. **J,K,** As in **A–G**, for n = 6 juvenile fish (15 dpf), whose eye diameter is approximately doubled^126^ relative to 7 dpf fish used above. Note the emergence of On-Off responses in the same IPL layers.

A very similar unmasking of latent On-Off terminals was also observed when following the pharmacological removal GABA_A_ and GABA_C_-receptors alone, leaving glycinergic signalling intact (Fig. 2H-I). By contrast, pharmacological blockage of either glycine, GABA_A_ or GABA_C_ in isolation had no such effects (Supplemental Figure S2A-F).

Together, these results show that approximately one third of larval zebrafish bipolar cells exhibit latent On-Off responses that are normally suppressed by GABAergic inhibition. Accordingly, polarity segregation at the level of bipolar cell output is shaped not only by feedforward mechanisms at the photoreceptor–bipolar synapse but also by inhibitory circuitry within the inner retina. Importantly, these effects were spatially structured (e.g. Fig. 2G), and reproducible across preparations (e.g. Fig. 2F), arguing against nonspecific network destabilisation.

### Outer retinal receptor systems permit intrinsic On-Off signalling

A likely origin of latent On-Off responses in bipolar cells (Fig. 2) are ‘mixed’ receptor systems that supply each polarity at the photoreceptor synapse (Fig. 1). If so, and if opposing polarity signals converge onto the same bipolar cells but are combined through nonlinear mechanisms, then the dominant polarity may mask the weaker one under baseline conditions. Because such masking may arise both from synaptic nonlinearities at the photoreceptor–bipolar synapse and from inhibitory circuits within the IPL, selectively disabling the receptors that supply one polarity should reveal pre-existing but suppressed responses of the opposite sign. Testing this prediction requires first consolidating our molecular understanding of how On and Off signalling is implemented in the zebrafish outer retina.

Zebrafish bipolar cells deviate from the canonical mammalian division in which AMPA and kainate receptors jointly define Off signalling, while mGluR6 mediates On signalling^69,73^ (Fig. 1B). Instead, Off bipolar cells predominantly express kainate receptors, whereas AMPA receptors are sparse. Next, mGluR6 expression is largely restricted to rod bipolar cells which are sparse in larvae^77,78^ which also lack functional rods^79,80^. Instead, EAATs, particularly EAAT5b, are broadly and robustly expressed across zebrafish bipolar cell types and have previously been implicated in On signalling^69,73^. This receptor landscape leads to two testable predictions. First, On signalling should rely primarily on EAATs rather than mGluR6. Second, Off signalling should depend almost entirely on kainate receptors. In both cases, suppressing one polarity should reveal enhanced signalling of the opposite polarity.

### EAATs, not mGluR6, mediate outer retinal On signalling in zebrafish

We first asked which receptor systems supply On signalling in the zebrafish retina. In line with transcriptomic predictions (Fig. 1B), pharmacological blockade of mGluR6 using AP4^81^ produced no reduction in On signalling; if anything, the fraction of On responses was slightly potentiated (Fig. 3A-E, Supplemental Movie M2, top row). This indicates that mGluR6 is not the primary mediator of outer retinal On signalling in larval zebrafish under the conditions examined here.

**Figure 3.**
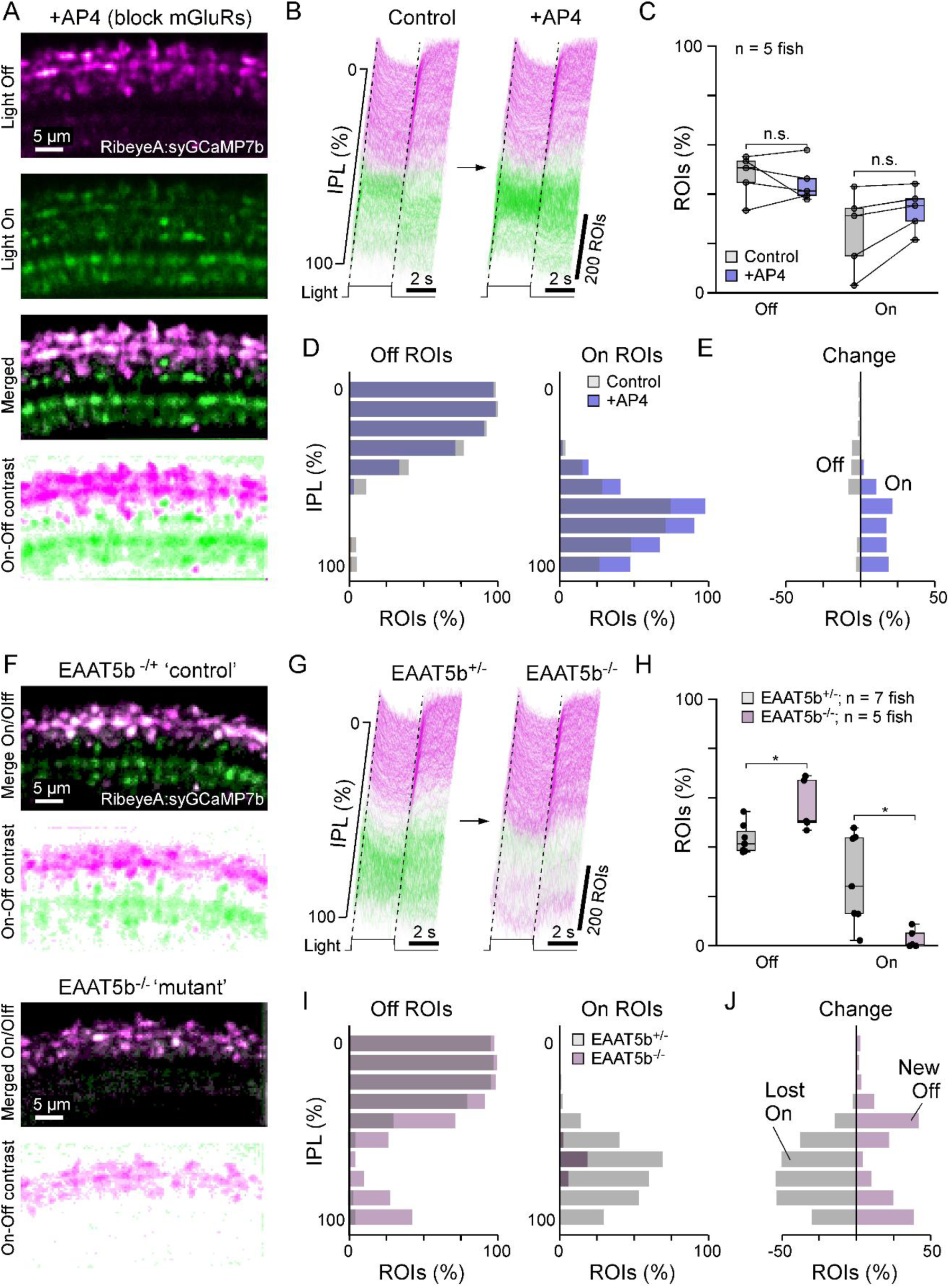
EAAT5b, not mGluR6, defines On pathway in larval zebrafish. **A–D,** As in Figure 2A–G, showing the effect of AP4-mediated block of group III glutamate receptors, which include mGluR6. This manipulation had minimal effects on the presence or distribution of On and Off responses in the larval zebrafish retina (n = 843 ROIs control, n = 993 ROIs drug, n = 5 fish). In **A** (bottom), On-Off contrast (Methods) is computed pixelwise across the example scan to illustrate the persistence of control-like On responses after treatment. **E,** Minimal change in the IPL distribution of On and Off responses between control and drug conditions, computed from the data in **D**. **F–J,** As in **A–E**, comparing EAAT5b^+/-^ zebrafish (“control”) with EAAT5b^-/-^ mutants. Unlike AP4 application, this manipulation robustly abolished On responses and unmasked latent Off responses (n = 1,238 ROIs “control” from 7 fish; n = 831 ROIs “mutant” from 5 fish). Statistical comparisons use Wilcoxon rank-sum tests (see Supplemental Table T2).

We next examined the role of EAATs, focusing on EAAT5b^82,83^. In EAAT5b homozygous mutants, On responses were nearly abolished, whereas heterozygous siblings showed no detectable deficit (Fig. 3F-I, Supplemental Movie M2, bottom left). Pharmacological blockade of EAATs yielded qualitatively similar but weaker effects, likely reflecting off-target effects, broader network perturbations and/or incomplete blockade (Supplemental Figure S3, Supplemental Movie M2, bottom right).

In all cases, suppression of On signalling was accompanied by potentiation of Off responses (e.g. Figs. 3I,J), specifically in the same IPL layers previously highlighted to comprise the latent On-Off terminals (Fig. 2G,I).

Together, these results demonstrate that EAAT5b is the dominant source of On signalling at the photoreceptor–bipolar synapse in larval zebrafish, and that removal of On-signalling unmasks a population of latent Off responses in the same IPL locations previously shown to comprise latent On-Off responses.

### Kainate receptors alone define Off signalling in zebrafish

We next asked which receptor systems supply Off signalling in the zebrafish retina. Guided by the transcriptomic data (Fig. 1B), we tested whether kainate receptors are sufficient to account for Off responses.

As predicted, pharmacological blockade of kainate receptors using ACET^84^ reliably abolished nearly all Off responses across the IPL (Fig. 4A–E), and simultaneously increased the fraction of terminals exhibiting On responses. These previously masked opposite-polarity responses were again confined to the same stereotyped IPL strata that contain the latent On-Off terminals (Fig. 4E; cf. Fig. 2G,I).

**Figure 4.**
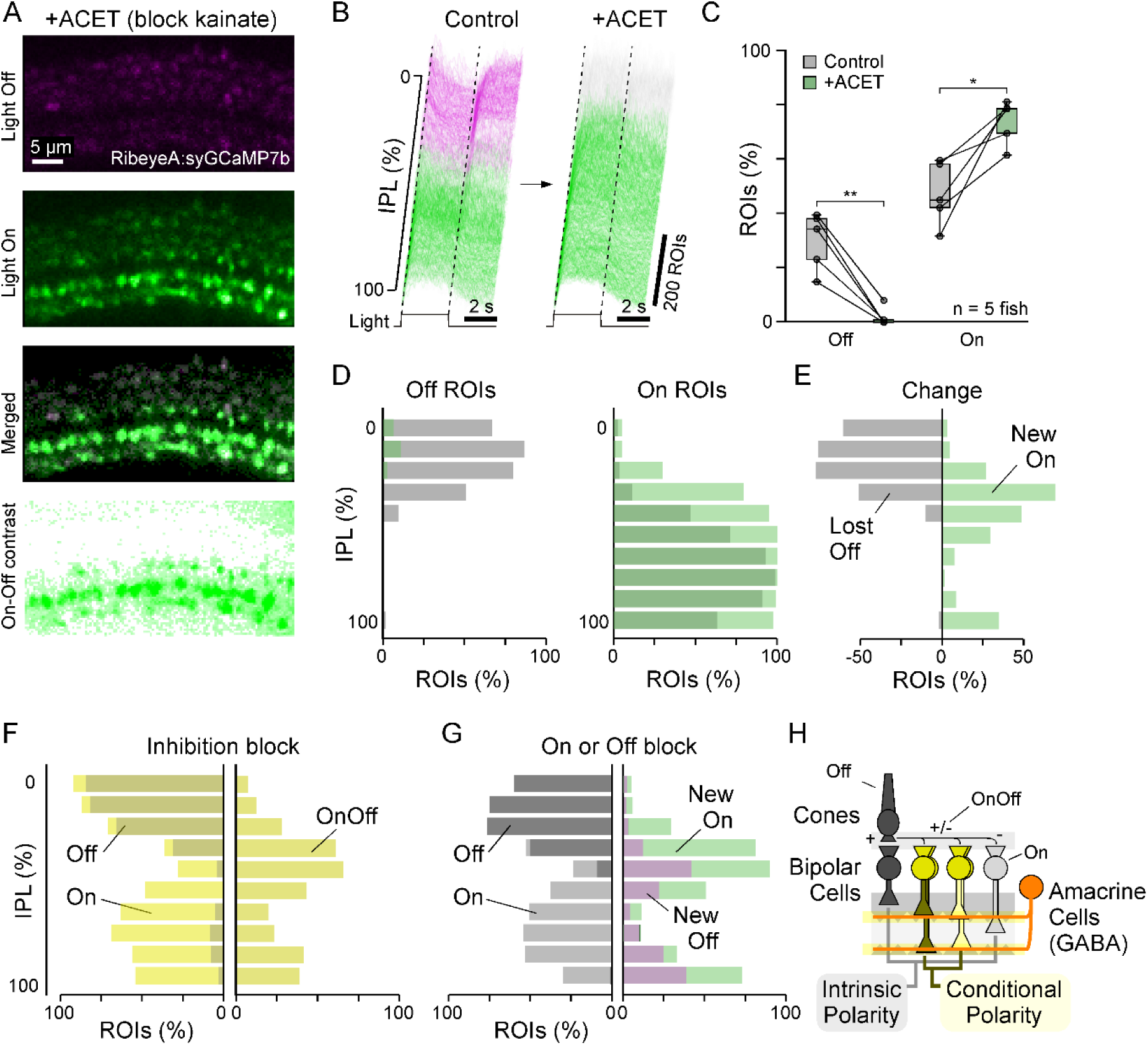
Kainate defines Off pathway in larval zebrafish. **A–E**, As in Figure 3A–E, showing the effect of blocking kainate receptors with ACET. This manipulation robustly abolished Off responses and unmasked latent On responses (n = 950 ROIs “control”, n = 920 ROIs “drug” from 5 fish). Statistical comparisons use Wilcoxon signed-rank tests (see Supplemental Table T2). **F,G,** Side-by-side comparison of IPL distributions of BC terminal responses unmasked by inhibition block (**F**; cf. Figure 2) and those unmasked by selective disruption of the On and Off pathways (**G**; Figures 3 and 4). Note that spatially equivalent BC populations are revealed in each case, suggesting common underlying bipolar cell types. In each figure, the left shows On and Off terminals superimposed on each other; right side represents the latent On-Off terminals after inhibition block (F) and newly emerged on (ACET application, green) or off (EAAT5b mutant, purple) terminals (G). **H,** Summary schematic of inferred latent On-Off bipolar-cell circuits in the larval zebrafish retina. Four types of bipolar cells (Off, 2x latent On-Off, and On) receive signal from cones, the latent On-Off bipolar cells may exhibit either on or off responses, determined by inhibition.

Together, these results demonstrate that Off signalling in larval zebrafish bipolar cells is driven almost exclusively by kainate receptors. Moreover, suppressing Off input unmasks latent On signalling in a defined subset of bipolar cell types. More broadly, these findings reinforce the conclusion that the apparent unipolarity of bipolar cell responses does not reflect an intrinsic constraint, but rather the active suppression of an opposing polarity supplied by distinct receptor systems (Fig. 4F–H).

This leads to a revised functional organisation of the larval zebrafish IPL. Rather than a strict division into On and Off layers, bipolar cell output can be partitioned into four functional strata starting from the inner nuclear layer (INL) side: Off, latent On-Off, On, and again latent On-Off. In this framework, the balance between On and Off signalling, including the effective position of the primary On/Off boundary, is therefore not hardwired, but actively maintained by inhibitory circuitry. This organisation may provide flexibility to adapt polarity coding to behavioural context, developmental stage, or retinal location (Fig. 4H).

### Inhibition enforces polarity segregation at the retinal output

The foregoing shows that inhibition within the IPL actively shapes the polarity of bipolar cell output. One synapse downstream, retinal ganglion cells (RGCs) receive both excitatory bipolar cell input and inhibitory input onto their dendrites. This raises a key question: do RGCs simply inherit the polarity imposed at the level of bipolar cells, or is polarity segregation actively re-established at this level?

To address this, we imaged dendritic activity of RGCs using an islet2b:TrpR; tUAS:mGCaMP6f line^57^ (Fig. 5A–E). Under control conditions, most dendrites exhibited predominantly On or Off responses, with a minority (∼15%) showing mixed On-Off responses, consistent with convergence of distinct bipolar cell inputs^16^.

**Figure 5.**
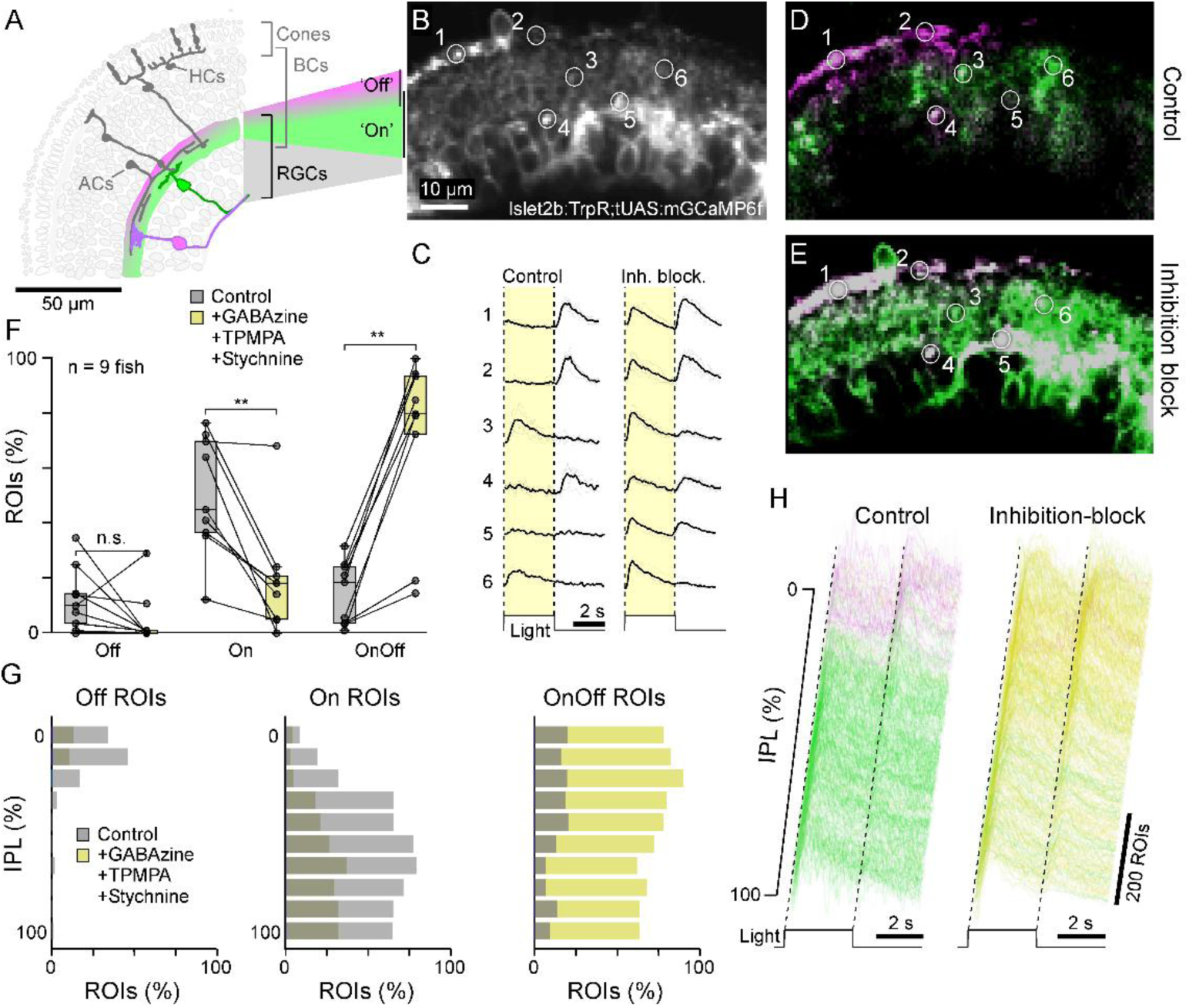
Inhibition block unmasks latent On-Off responses in retinal ganglion cells. **A–H**, As above, but now showing light responses of retinal ganglion cell (RGC) dendrites and somata in the larval zebrafish eye (Islet2b:mGCaMP6f) ^57^ under control conditions and following pharmacological block of GABAA, GABAC, and glycine receptors. At this stage of retinal processing, 74% of ROIs exhibited On-Off responses after disinhibition, up from 15% in control conditions (n=889 ROIs “control” and n=1,153 ROIs “drug” from 7 fish). Statistical comparisons use paired t-test (see Supplemental Table T2).

Blocking GABA and glycinergic inhibitory transmission dramatically altered this pattern. Following disinhibition of both, the vast majority (>80%) of RGC dendrites exhibited robust On-Off responses (Fig. 5B-H, Supplemental Movie M3). This effect was substantially stronger than observed in bipolar cells (cf. Fig. 2) and, critically, was not restricted to specific IPL strata (Fig. 5G), indicating a general role for inhibition in enforcing polarity segregation at this next level of retinal processing.

These results demonstrate that polarity segregation in RGC dendrites is not simply inherited from upstream circuits but is actively maintained by inhibitory mechanisms. The pronounced increase in mixed-polarity responses upon disinhibition, including in IPL layers where the polarity of bipolar cells remained ‘stable’ (cf. Fig. 4F-H), further indicates that a second, functionally distinct layer of inhibition operates at the level of RGCs to constrain polarity. The consistency of this effect across cells and recording sites suggests a systematic circuit mechanism rather than a global change in excitability.

### Polarity segregation is again re-established in central visual circuits

If polarity is actively sculpted within the retina (Figs. 1-5), an important question is whether this principle extends to downstream visual circuits. Retinal ganglion cells (RGCs) project to multiple arborisation fields (AFs) in the brain in a type-specific manner^85^. This provides an opportunity to examine how polarity is represented across distinct sets of RGC types, and their target regions.

We focused on AF10 (the optic tectum), the largest and functionally most general visual projection area, and AF9, a smaller target associated with more specialised processing^86^. Using the same reporter line as for retinal imaging, we recorded activity in RGC axon terminals within these regions (Fig. 6A-C).

**Figure 6.**
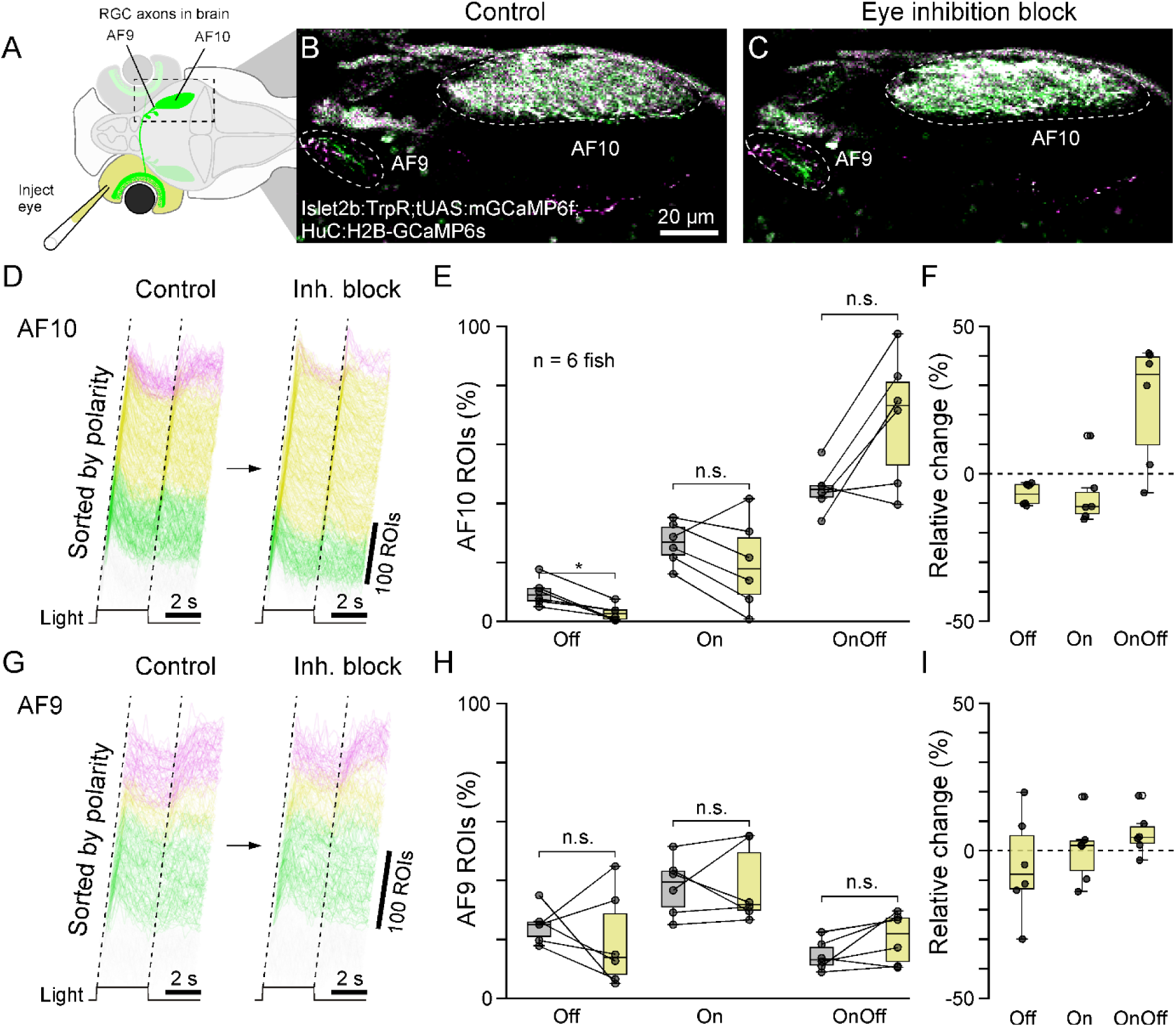
Disinhibition of the eye alone causes limited disruption of polarity coding in RGC projections. **A–I**, As in Figure 5, but now imaging RGC axon terminals in the brain. Unlike their dendritic and somatic counterparts in the retina, RGC axon terminals were comparatively resistant to disruption of inhibitory circuitry in the eye, implying an additional level of local control in the brain. Two major projection targets were analysed separately: AF10, the principal target of most RGC axons (**D–F**), and AF9, the largest pretectal projection field innervated by a smaller subset of RGC types (AF9: n = 281 ROIs “control” and n = 283 ROIs “drug”; AF10: n = 596 ROIs “control” and 1081 ROIs “drug”, from 6 fish, **G–I**). Statistical comparisons use Wilcoxon signed rank test (6E) and paired t-test (6H) (see Supplemental Table T2).

The strong shift toward On-Off responses observed in the retina following disinhibition (Fig. 5) predicts a comparable increase in mixed-polarity signalling in these central projection fields. However, this was not the case. In AF10, the fraction of On-Off responses increased only moderately (from ∼45% to ∼75%), representing a substantially weaker relative effect than in the retina (Fig. 6D-F). Moreover, in AF9, RGC axons exhibited no significant change towards On-Off signalling (Fig. 6G-I).

These observations indicate that, despite originating from the same RGC population, polarity is again re-segregated in central targets, likely via presynaptic inhibition of RGC axon terminals in the brain.

### Further levels of inhibitory polarity control in central neurons

RGC outputs provide input to central neurons, many of which reside in the optic tectum^86^, a structure highly enriched in inhibitory interneurons^87^. We therefore asked how polarity is represented at the level of postsynaptic visual brain neurons and whether it remains subject to additional levels of inhibitory control.

For this, we simultaneously imaged RGC axonal terminals and central neurons by combining an islet2b:TrpR; tUAS:mGCaMP6f line with a pan-neuronal reporter (HuC:H2B-GCaMP6s) (Fig. 7A-C). Because these reporters differ both spatially and in signal structure (diffuse neuropil versus nuclear-localised signals), the two populations could be readily separated. Under control conditions, RGC terminals in the brain exhibited response profiles like those observed previously (Fig. 6). By contrast, and in line with previous descriptions (e.g. Ref^74^), central neurons showed predominantly unipolar responses: Many (>50%) were unresponsive to our widefield stimulus, followed by On- (∼35%) and Off-responsive neurons (∼10%). Only a small minority (∼1%) exhibited On-Off responses (Fig. 7D,E).

**Figure 7.**
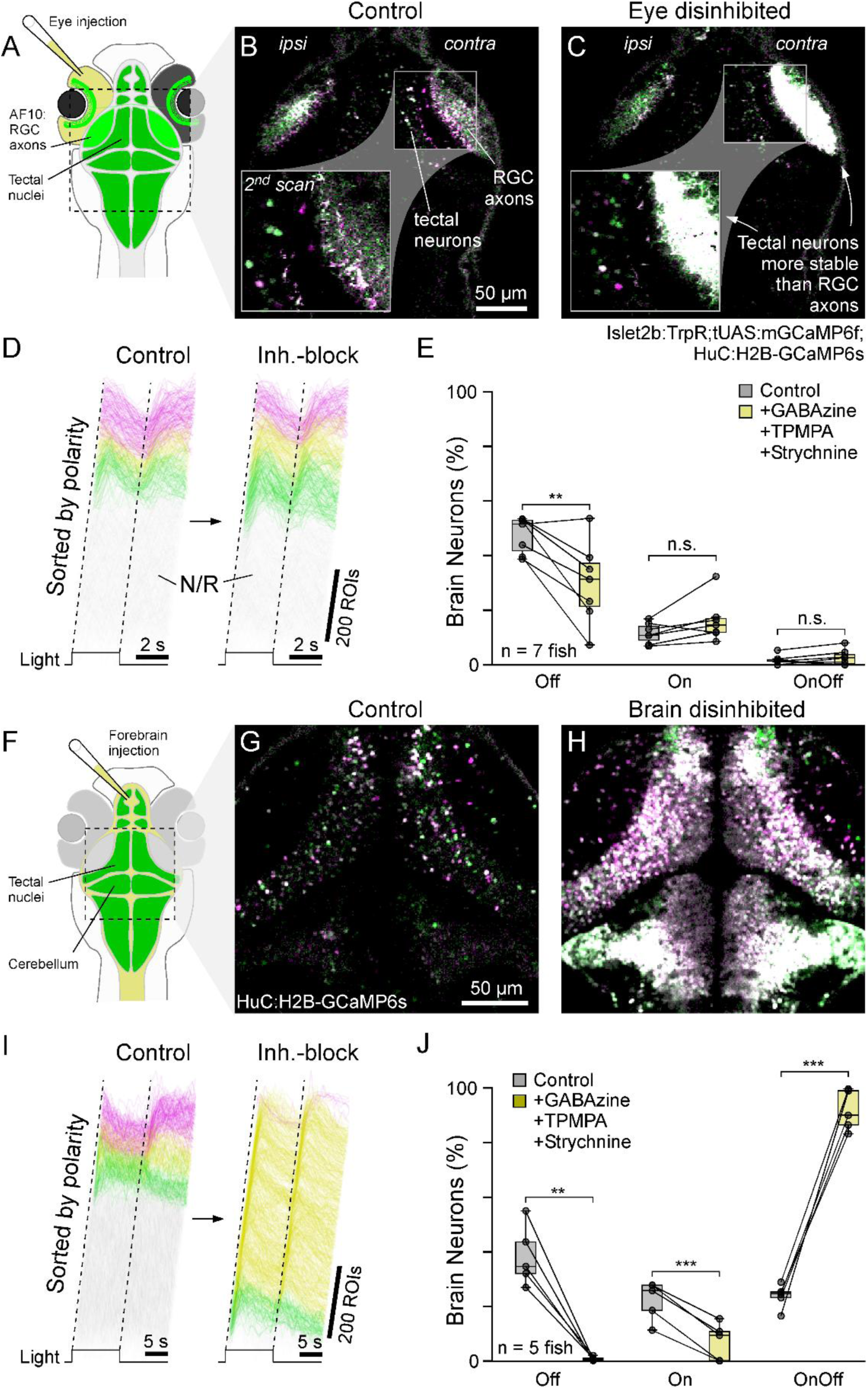
Disinhibition of the brain, but not the eye, unmasks widespread latent On-Off responses in central visual neurons. **A–E**, As in Figures 5 and 6, but now simultaneously imaging RGC projections and local brain neurons in the tectum and downstream areas using a double-transgenic line (islet2b:TrpR; tUAS:mGCaMP6f; HuC:H2B-GCaMP6s). Disinhibition of retinal circuits had only minor effects on the distribution of response polarities in the brain. **F–J,** By contrast, local pharmacological inhibition block in the brain caused nearly all central neurons to exhibit robust On-Off responses. For eye disinhibition experiments, n = 1,833 ROIs “control” and n = 2,131 ROIs “drug”, from 7 fish. For brain disinhibition, n = 4,533 ROIs “control” and n = 6,175 ROIs “drug”, from 5 fish. Statistical comparisons use paired t-tests (see Supplemental Table T2).

Disinhibition of the retina led to a moderate inflation of mixed-polarity responses in RGC terminals within AF10 (Fig. 7C), in line with the foregoing (Fig. 6). However, at the same time, the polarity of central neurons remained largely unaffected (Fig. 7C-E). This indicates that mixed-polarity signals present at the level of retinal output are again filtered or reshaped before being expressed in central neuronal activity.

To test whether this reflects an additional layer of local inhibitory control, we therefore directly blocked both GABA and glycinergic inhibition within the brain (Methods, Fig. 7F-J, Supplemental Movie M4). This manipulation produced a dramatic shift: now nearly all central neurons exhibited strong On-Off responses. A similar effect was observed when only blocking GABAergic transmission (Supplemental Figure S4). In both cases, the consistency of these transformations argues against nonspecific network effects.

Taken together, these results reveal a consistent principle across the visual system of the zebrafish. At each stage, from BCs (Fig. 1-4) to RGCs (Fig. 5), their axon terminals in the brain (Fig. 6), and to central neurons (Fig. 7), signals capable of encoding both On and Off polarities are present across the stages examined. However, at each stage, inhibitory circuits selectively suppress one component to generate apparently segregated responses.

Thus, polarity segregation is not only inherited as a fixed property from earlier processing stages, but also repeatedly reconstructed through the interaction of excitatory inputs and local inhibitory mechanisms.

### Dynamical switching between mixed and unipolar responses depending on light levels

A circuit architecture in which mixed-polarity signals are available but actively suppressed predicts that polarity coding should be dynamically adjustable. For example, the balance between mixed and segregated responses may depend on the effective signal-to-noise^35^: as photon catch increases, either through eye growth or higher ambient light levels, visual circuits may be able to tolerate, or exploit, mixed-polarity signalling more readily. Conversely, under lower light levels, stronger polarity segregation may help stabilise visual representations.

We first tested this idea developmentally by imaging bipolar cell terminals in juvenile (15 *dpf*) zebrafish, when eye diameter is approximately doubled relative to 6–8 *dpf* larvae. Because photon catch scales approximately with aperture area, this corresponds to an estimated fourfold increase in light capture. Strikingly, unlike larvae, juvenile bipolar cells already displayed a small but robust population of mixed On-Off responses under control conditions (Fig. 8A-F). These responses closely resembled the latent On-Off terminals unmasked in larvae by disinhibition, including their stereotyped IPL distribution (Fig. 2A-G). Thus, bipolar cell types that are normally held in an apparently unipolar state in larvae can express mixed polarity without pharmacological manipulation at later developmental stages.

**Figure 8.**
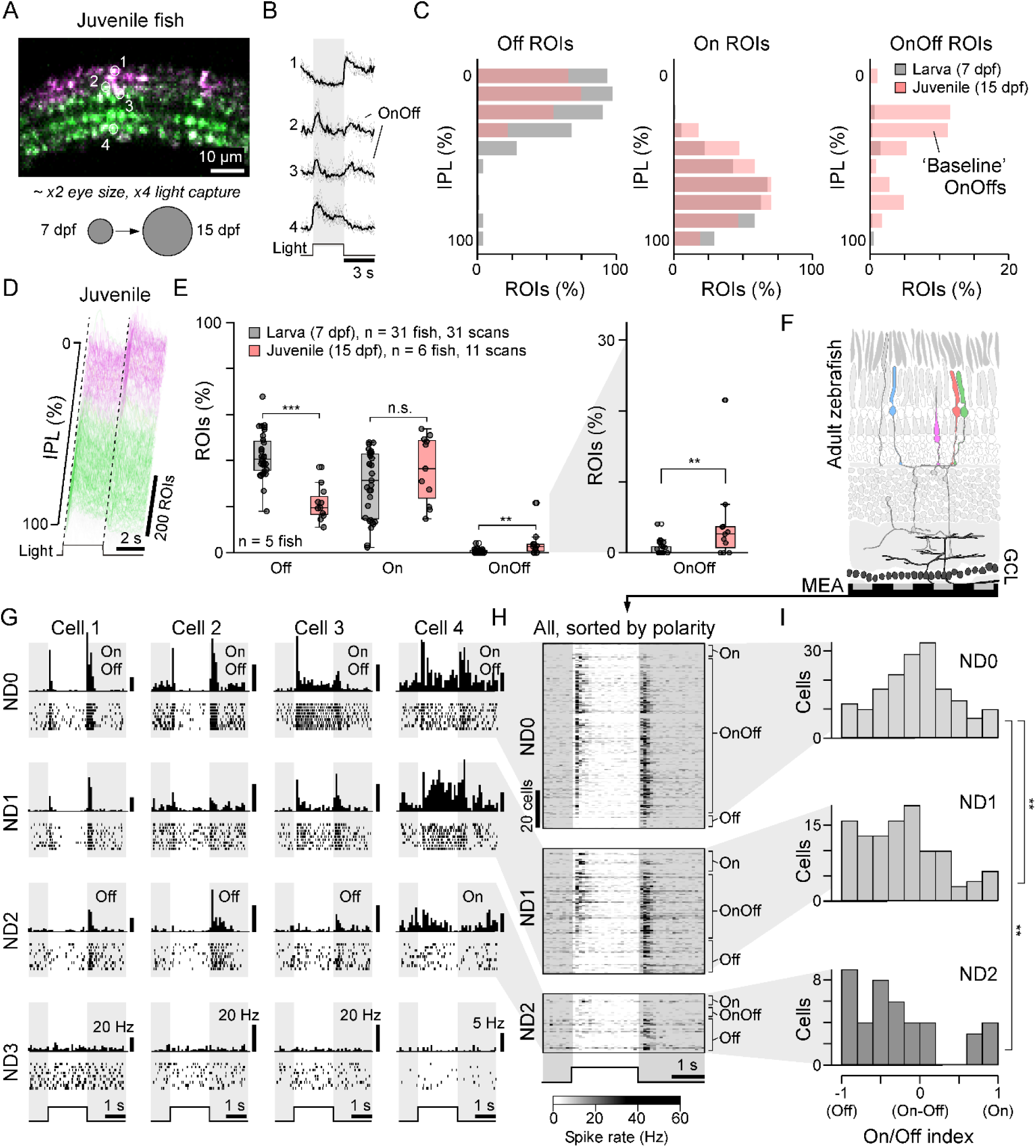
Polarity coding switches dynamically with effective light level. **A–E**, Bipolar cell terminal responses in juvenile zebrafish at 15 dpf. Unlike 6–8 dpf larvae, juveniles exhibited a small but robust population of mixed On-Off responses under control conditions, concentrated in IPL strata matching the latent On-Off terminals revealed by disinhibition in larvae. Statistical comparisons use Wilcoxon rank-sum tests as indicated in Supplemental Table T2. **F-I**, Multielectrode-array recordings from the adult zebrafish ganglion cell layer (F) across light levels (G-I). Shown are 4 example cells (G), an overview of all responsive units per light level (H) and their corresponding distribution of OnOff index (I; with OnOff-index -1 to 1 corresponding to ‘perfect’ Off and On, respectively, with OnOff responses occupying intermediate values;). n = 2 retinas yielded 562 spike-sorted units, of which 170 (ND0), 110 (ND1), 42 (ND2) and 0 (ND3) passed a response quality criterion (Methods). A ‘white’ contrast series (100-10% in 10% steps) was presented, but for simplicity only the responses to the 90% contrast step were used for analysis (H,I). For improved visualisation, individual cells shown in (G) combine the responses to the 90, 80 and 70% contrast steps, which were qualitatively similar (Methods). Kruskal-Wallis test and post-hoc pairwise Mann-Whitney U tests were used for statistical comparison (see Supplemental Table T2).

We next asked whether this developmental shift continues into adulthood and whether polarity coding can also switch dynamically within the same visual system as a function of light level. Because adult zebrafish eyes are too large and pigmented for routine *in vivo* two-photon access to the retina, we used multielectrode-array recordings from the ganglion cell layer (Fig. 8F, Methods), which is dominated by retinal ganglion cells. Strikingly, under photopic conditions, most visually responsive adult units exhibited robust mixed On-Off spiking responses (Fig. 8G-I; Neutral Density Filter 0, ND0). Thus, the developmental emergence of mixed-polarity signalling observed in juvenile bipolar cells is not transient but appears to progress further in the adult retinal output, where mixed On-Off coding becomes the dominant retinal output mode under bright conditions. However, when the same units were challenged with progressively lower light levels (ND1 and 2, corresponding to 10- and 100-fold brightness decrease, respectively), their responses shifted toward a more segregated regime, becoming increasingly unipolar and Off-dominated (Fig. 8G-I, middle/bottom), before ceding completely (ND3, 1000-fold decrease).

Together, these experiments show that mixed On-Off coding is not merely revealed by pharmacological perturbation but emerges naturally as effective light input increases across development and into adulthood. Conversely, reducing light level or contrast favours more segregated, polarity-biased signalling. Thus, polarity organisation is dynamically regulated rather than fixed, consistent with the idea that inhibition controls access to a latent mixed-polarity mode according to the prevailing visual regime.

### Latent On-Off coding is conserved in mammals

To test whether the principle of latent On-Off responses generalises beyond zebrafish, we examined polarity coding in the rod-dominated the mouse retina (Fig. 9). Using *ex vivo* recordings^26^ from the ganglion cell layer (which comprises RGCs and displaced amacrine cells), we quantified the proportions of On, Off, and On-Off responses under control conditions and following pharmacological disinhibition of GABAergic transmission.

**Figure 9.**
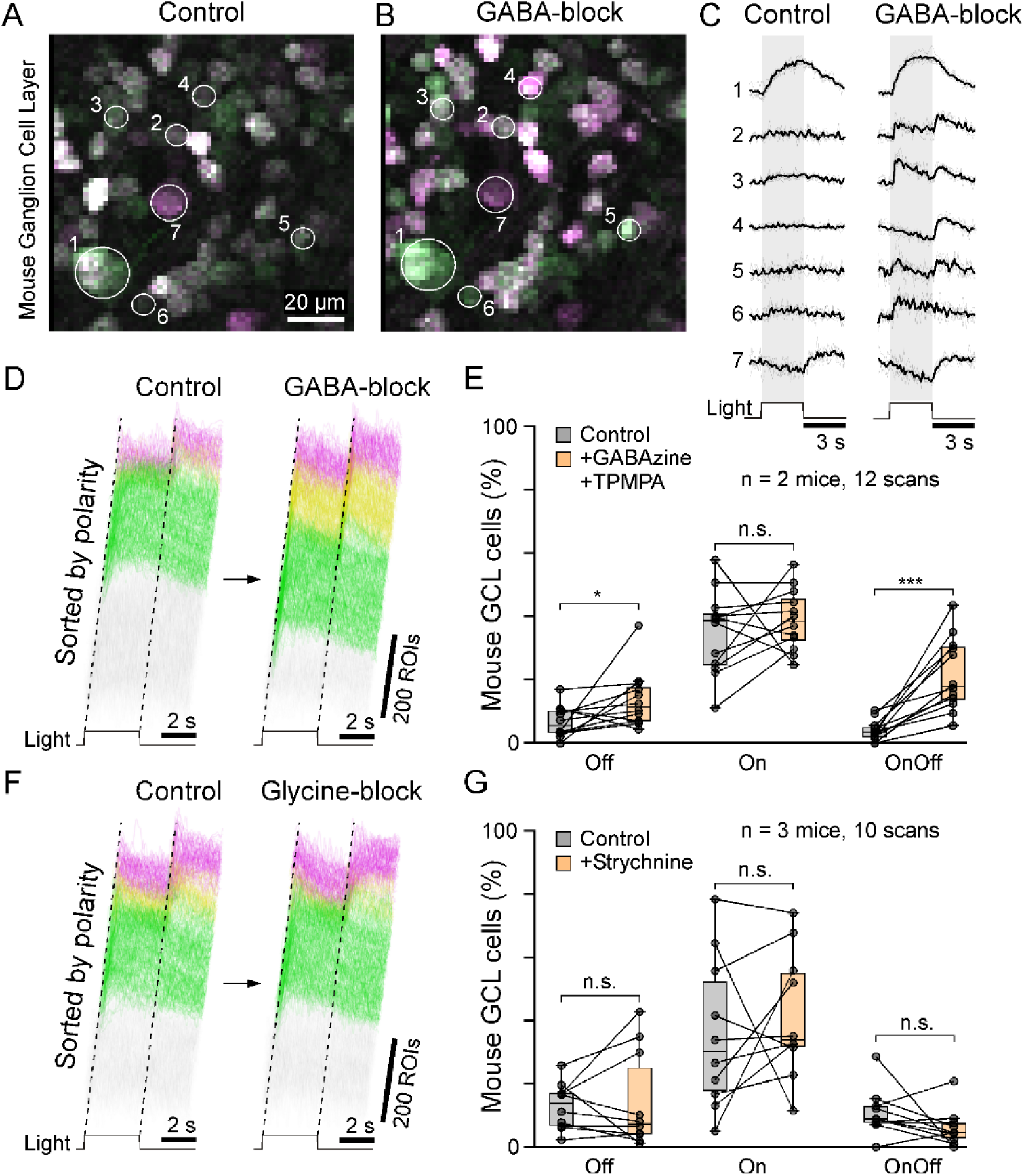
GABA-block unmasks latent On-Off responses in mouse retinal ganglion cells. As in Figure 5, but now comparing RGC soma responses in mouse retinal wholemounts following bulk electroporation^26,127^ of the calcium indicator Oregon-Green 488 BAPTA-1 (OGB-1) under control conditions and after bath application of TPMPA and GABAzine to block GABAergic neurotransmission. As in zebrafish, this manipulation robustly unmasks latent On-Off responses in 20% of RGCs, up from 4%; n = 12 scans from 2 mice. **F,G,** By contrast, blocking glycinergic transmission had no such effect (data reanalysed from Ref^88^). n = 10 scans from 3 mice. Statistical comparisons use paired t-tests (see Supplemental Table T2).

Consistent with previous reports^26,27^, most RGCs in control conditions exhibited predominantly On or Off responses, with a smaller fraction showing mixed polarity. Blocking GABAergic transmission (TPMPA plus GABAzine) then led to a marked increase in On-Off responses across the population (Fig. 9A-E).

By contrast, blocking glycinergic transmission (strychnine) ^88^ did not unmask additional On-Off responses (Fig. 9F-G), in line with foregoing observations from zebrafish.

Together, these results suggest that latent On-Off coding and its regulation by GABAergic inhibition are features likely conserved across vertebrates, rather than specialisations of zebrafish.

## DISCUSSION

### Polarity segregation as a reusable circuit operation

Our results support a shift from a mainly feedforward view of On versus Off pathway splitting to a distributed one. Rather than being established once at the photoreceptor–bipolar synapse and subsequently propagated, polarity segregation emerges as a circuit state that is repeatedly constructed at multiple stages of processing. At each level examined, from bipolar cells (Figs. 1-4) to retinal ganglion cells (Figs. 5,6,8,9) and central visual neurons (Fig. 7), signals capable of encoding both polarities are widely present, but inhibitory circuitry determines which component is expressed.

This reframes how mixed responses may be interpreted. In the visual system of larval zebrafish at a minimum, and plausibly more broadly across species and modalities, mixed-polarity signalling appears to be an intrinsic property of the circuit rather than reflecting crosstalk between otherwise segregated pathways. In this view, unipolar responses arise when one component is selectively suppressed, making polarity a dynamically regulated outcome that circuits can repeatedly apply.

### Functional advantages of latent mixed-polarity signalling

A key implication of this framework is that maintaining access to both polarities may enhance functional flexibility. In this view, visual circuits retain latent mixed-polarity signalling, allowing polarity to be dynamically reweighted in response to changes in computational demands, for example those associated with light levels^35,50^, stimulus statistics^89–92^, or behavioural state^93–95^. The ability to dynamically switch between mixed and segregated polarity codes, as demonstrated here, provides a concrete mechanism by which such flexibility can be implemented without requiring structural circuit changes.

#### Signal-to-noise and luminance regime

A primary driver of polarity organisation is likely signal-to-noise ratio. Under low-light conditions, stronger polarity segregation can help stabilise representations by limiting ambiguous signals^11,44^. This is consistent with the prominence of clean On and Off channels in rod-dominated species, including many mammals. At the mechanistic level, receptor systems associated with these pathways, including the mGluR6 cascade^96^ and, potentially, the high peak currents associated with AMPA receptors^97^, are well suited to detecting and amplifying low-amplitude inputs, which may favour their deployment under dim-light conditions. Conversely, receptor systems such as kainate receptors and EAATs, which are more prominent in cone-dominated and diurnal systems in our comparative analysis (Fig. 1), may be better suited to operating in higher signal regimes, where input amplitudes are larger and the constraints imposed by noise are reduced. This is because their comparatively lower gain and slower kinetics may favour graded, sustained signalling over the rapid amplification required for detecting weak inputs.

Our results directly support this view. In zebrafish, mixed-polarity signalling emerges progressively with increasing effective light input (Fig. 8): it is largely latent in larvae, becomes partially expressed in juveniles, and dominates retinal output in adults under photopic conditions. Within the same adult retina, this balance can be rapidly reconfigured, with lower light levels or reduced contrast shifting responses toward a more segregated, predominantly unipolar regime. Thus, polarity organisation tracks the effective signal regime both across development and on fast timescales.

In addition, polarity balance can vary over circadian timescales: zebrafish show a rebalancing of On and Off signals across the day^98,99^, consistent with ongoing adjustment to changing luminance conditions. Together, these observations suggest that polarity organisation adapts to the effective noise regime rather than remaining fixed.

#### Adaptation to environmental and behavioural statistics

Polarity balance is also shaped by the contrast structure of behaviourally relevant stimuli. Some tasks favour emphasising one polarity over the other, depending on whether relevant signals are predominantly increments or decrements. In zebrafish, the prey-capture acute zone^91^ shows a strong On-bias^57,100^, consistent with detecting UV-bright prey against a darker background^89^, whereas other retinal regions are more balanced. In mice, retinal physiology likewise varies systematically with retinal position^92,101^, including differences in the relative representation of On and Off pathways^102–104^, consistent with regional specialisation for distinct visual environments^90^. Such examples indicate that polarity coding is tuned not only by noise constraints but also by task-specific stimulus statistics. Latent mixed-polarity signalling provides a flexible substrate from which local circuits can derive these specialisations without requiring rigidly separate pathways.

#### Behavioural state and dynamic task switching

Visual demands can also change rapidly with behavioural state^93,94^. Zebrafish, for example, readily switch between exploratory behaviour and localised prey capture^95^, which likely place different demands on visual processing. Exploration may benefit from balanced access to both On and Off signals for scene interpretation and navigation, whereas prey capture favours On-biased channels tuned to bright targets^89^. Latent mixed-polarity circuitry would allow the system to reweight existing channels accordingly. More generally, this also aligns with the emerging view that ancestral cone circuits are antagonistically organised to flexibly extract behaviourally relevant signals at the network level^68,74^, with polarity balance adjusted as task demands change.

### Evolutionary considerations

From an evolutionary perspective, a parsimonious interpretation is that mixed-polarity signalling reflects the ancestral condition, with strongly segregated On and Off channels emerging as a derived specialisation in some lineages. The ancestral vertebrate visual system is thought to have operated under predominantly diurnal, light-rich conditions^63,105–107^, favouring high information throughput over strict noise minimisation. In this regime, mixed-polarity representations may have been advantageous as a flexible and information-rich coding strategy^52^. Consistent with this view, non-mammalian vertebrates often exhibit widespread mixed-polarity signalling^52–57^, and the robust presence of ‘all four’ receptor-variants (Fig. 1A) in lamprey bipolar cells (Fig. 1B) strongly suggests they were fully available for use in the ancestral vertebrate eye.

In contrast, around a quarter billion years later, early mammals experienced a prolonged nocturnal bottleneck^108^, during which visual systems were optimised for low-light conditions. Under these constraints, stronger and earlier polarity segregation may reduce noise and limit the propagation of ambiguous signals. And yet, mixed responses can still be increasingly deployed by mammalian retinas under appropriate conditions^35,50^, indicating that some degree of latent On-Off capability is retained.

A related perspective comes from the evolutionary origins of bipolar cell pathways. Recent work^109^ indicates that the ‘original’ cone-bipolar channels were likely Off, whereas the ancestral primary rod bipolar pathway^69,110^ is, and remains^111^, On. Within this framework, On-cone bipolar cells emerged later through the recruitment of On-signalling mechanisms onto an existing Off-cone bipolar scaffold^109^. If so, early On-cone bipolar cells would have at least transiently exhibited mixed On-Off signalling before becoming selectively On. Our findings are consistent with the idea that such mixed states are not merely transitional but may persist as a latent capability in many bipolar cell types.

More broadly, key elements of visual polarity splitting extend beyond vertebrates. In insects, for example, On and Off pathways are also segregated from the first synapse of vision^3,112^, and disinhibition can enhance mixed On-Off responses^4,113^, paralleling our observations. Other visually sophisticated protostomes, such as cephalopods, likewise exhibit On-Off pathway organisation in the optic lobe^114,115^, although the underlying circuit mechanisms and neurotransmitters remain less well resolved^116^. Together, these observations open the possibility that flexible, inhibition-dependent control of polarity may be a widely shared feature of visual systems, with strict pathway segregation representing a derived and context-dependent refinement.

### GABA versus glycine

In vertebrate visual circuits, most inhibition arises from the interplay of GABAergic and glycinergic neurotransmission^46,117^. In both retina and brain, our results point to a prominent role of GABAergic rather than glycinergic inhibition in shaping response polarity. This observation is broadly consistent with known organisational features of retinal circuitry. The distinction between GABAergic and glycinergic amacrine cell types is ancient^118^ and strongly correlated to distinct anatomical and functional properties: GABAergic circuits largely operate laterally within IPL strata, while glycinergic circuits often span layers vertically and are implicated in crossover interactions between On and Off pathways^119^. Such crossover inhibition^25^, Such crossover inhibition, commonly associated with glycine, is mechanistically distinct from the polarity control described here: classical crossover inhibition acts between already segregated On and Off pathways, whereas our data suggest a mechanism that gates the expression of mixed-polarity signals before or within individual output channels. The weak effect of glycine block is therefore intriguing and suggests that polarity selection is mediated by a separable GABAergic circuit motif rather than by canonical crossover inhibition alone. Together, these considerations reaffirm that in the retina, GABAergic and glycinergic inhibition implement partially distinct computational roles, with GABAergic circuits contributing more directly to polarity selection under the conditions examined here. Whether a similar division of labour extends to central visual circuits, where inhibitory functions are less well characterised, remains an open question.

### Limitations and future directions

First, our functional analyses focus primarily on larval zebrafish. While this system provides access to a cone-dominated and perhaps an ancestrally informative^68^ vertebrate retina, the extent to which the principles described here generalise across developmental stages, species, and visual ecologies remains to be established. In particular, testing across a broader phylogenetic range, including species with different evolutionary histories and luminance niches^120,121^, will be important to assess how universal latent mixed-polarity signalling is.

Second, pharmacological manipulations may introduce network-level effects. However, the convergence of pharmacological and genetic approaches, together with the spatial specificity of observed changes, supports the interpretation that these results reflect unmasked circuit capacity rather than nonspecific destabilisation. Nevertheless, future work using cell-type-specific perturbations will be important to dissect underlying mechanisms.

Third, transcriptomic co-expression does not necessarily translate to functional receptor contributions^69,71^. Synaptic localisation, receptor density, and circuit context will influence effective polarity.

Fourth, comparative analyses remain limited by available datasets, which remain biased toward mammals (Fig. 1). In addition, species with domesticated (pig, sheep, cow, chicken) or commensal lifestyles (house mice), and humans themselves, all may experience atypical light environments, potentially decoupling present-day ecological conditions from evolutionary history.

## Conclusions

Together, our results support a revised view of On-Off pathway splitting. Rather than a fixed transformation established at a single synapse, polarity segregation may be better understood as a dynamic and distributed computation that is repeatedly constructed through the interaction of excitation and inhibition.

In this framework, mixed-polarity signalling represents a latent operating regime that can be flexibly recruited, while clean On and Off channels emerge as a regulated outcome of suppression. This perspective reconciles variability across species and conditions and suggests that canonical neural computations arise not only from fixed circuit motifs, but from flexible operations whose expression depends on the prevailing signal regime.

## SUPPLEMENTAL MOVIES

**Supplemental Movie M1.** Averaged recording movies from zebrafish bipolar cell terminals when presented white full field stimulation (3x real time). Two conditions were shown in this video (7 dpf control, 7dpf GABA+glycine block,) as indicated at the bottom of each half. The white stimulus presented is indicated at the top left corner for each condition.

**Supplemental Movie M2.** As M1, here showing four different conditions for on-pathway blockage: control, AP4-application, EAAT5b^-/-^ mutant, TBOA application (3x real time).

**Supplemental Movie M3.** As M1, here showing RGC dendrites in control and inhibition block condition (3x real time).

**Supplemental Movie M4.** As M1, here showing brain neurons after local disinhibition in the brain. (5x real time).

## SUPPLEMENTAL TABLES

**Supplemental Table T1.** Glutamate receptor / transporter genes used for comparative transcriptomics analysis (Fig. 1)

**Supplemental Table T2.** Details on all statistical tests used and their p-values**. Supplemental Table T3.** Spiking circuit parameter list, as used for MEA spike sorting.

## METHODS

### Animals

*Zebrafish*. All experimental procedures were conducted in compliance with the UK Animals (Scientific Procedures) Act 1986 and carried out under UK Home Office guidelines, with ethical approval granted by the Animal Welfare Committee at the University of Sussex. Zebrafish larvae (*Danio rerio*) were maintained in petri dishes at a density not exceeding 50 animals per dish, at a controlled temperature of 28°C and under a 14:10 hour light/dark cycle (lights on at 08:00, lights off at 22:00), with 15-minute gradual luminance transitions at both dawn and dusk to simulate natural photoperiod conditions. From 1 day post-fertilisation (dpf), larvae were raised in fresh fish facility system water supplemented with 200 μM 1-phenyl-2-thiourea (PTU; Sigma-Aldrich, catalogue no. P7629) to inhibit melanogenesis throughout the experimental period^122^ (Karlsson, Von Hofsten and Olsson, 2001). Adult zebrafish were housed in ZebTec Active Blue (TECNIPLAST) system, with temperatures 26.5 °C and a 14:10 hour light/dark cycle. The following published transgenic lines of zebrafish were used: *Tg(ribeyeA:syGCaMP8m)*, *Tg(ribeyeA:syGCaMP7b)*, *Tg(isl2b:TrpR; tUAS:mGCaMP6f)*, *Tg(HuC:H2B-GCamP6s), eaat5b^-/-^*knock-out^83^.

*Mouse*. All two-photon Ca²⁺ imaging experiments were performed at the University of Tübingen in accordance with German animal experimentation laws and approved by the institutional animal welfare committee. We used retinae (n=4 for GABA-block experiments, n=6 for Glycine-block experiments) from C57Bl/6J mice (n = 2 for GABA-block experiments, n = 3 for Glycine-block experiments; JAX 000664) of either sex, aged 4–16 weeks, housed under a 12h/12h light/dark cycle at 22°C and 55% humidity.

### Tissue Preparation

#### Zebrafish imaging

Zebrafish larvae were immobilised in 2% low melting point agarose (Fisher Scientific), for retinal recordings, larvae were mounted laterally with the eye of interest oriented upward toward the objective; for recordings of the arborisation field (AF) and optic tectum, larvae were mounted dorsally. The imaging chamber was filled with E2 fish medium, submerging the agarose-fixed larvae. Eye movements were further prevented by injection of α-bungarotoxin (1 nL of 2 mg/ml; Tocris) into the ocular muscles behind the eye.

#### Zebrafish multielectrode array recordings

One litre of ringer solution (255 mOsm/L) was freshly prepared for each experiment. To one litre of purified water (18.2 MΩ·cm; Purelab Chorus, ELGA), the following solutes were added (in mol/L): NaCl (0.0967), KCl (0.00201), MgCl_2_6·H_2_O (0.000787), NaH_2_PO_4_ (0.00100), Glucose (0.0154), and NaHCO_3_ (0.0201). The solution was bubbled with carboxygen (95% O_2_ / 5% CO_2_) for at least five minutes, before adding CaCl_2_ (0.00207 mol/L) and L-glutamine (0.000411 mol/L). Zebrafish adults were dark adapted for at least 12 h overnight. Under dim red light zebrafish were culled via immersion in 1-2°C water for ∼30s, followed by decapitation. The head was placed in ringer solution. Using curved scissors (FST 15000-04) the eye was lifted and the optic nerve cut. Following enucleation, a needle (30 gauge, 13mm, unisharp) was used to make a hole in the anterior of the eye above the lens. Using forceps (MSS 11200-14) and curved scissors the cornea and lens were removed. Vitreous was removed from the eyecup using forceps. The sclera and choroid were removed from the retina by placing forceps between the retinal pigment epithelial (RPE) and the choroid, leaving as much RPE attached to the retina as possible. The inner limiting membrane was removed from the inside of the retinal cup using forceps. Four incisions using straight edged scissors (FST 15004-08) were made from the retinal edge approximately one-third of the way towards the optic nerve. The retina was flattened onto filter paper (MCE Membrane Filter, AABG0100), with the photoreceptor layer against the filter paper. The filter paper was briefly (∼5 s) transferred onto paper towel to remove solution, facilitating the flattening of the retina. The filter paper and retina were transferred to the MEA chamber (which had been filled with ringer solution) and placed onto the electrode array, with the ganglion cell layer in contact with the electrode array. Ringer solution was removed from the MEA chamber using a pipette, to aid attachment of the retinal tissue to the electrode array. Fresh ringer solution was then pipetted onto the filter paper and tissue to fill the chamber and to consolidate retinal attachment to the electrode array. A harp slice grid (ALA HSG-MEA-5B, 0.4g) was placed on top of the filter paper. The MEA chamber was connected to the headstage and the retina left to dark adapt for 45 minutes. The tissue was perfused throughout the experiment with ringer solution, which was continuously bubbled with carboxygen (95% O_2_ / 5% CO_2_).

#### Mouse

All procedures were performed under dim red light (>650 nm). Animals were dark-adapted for >1h, anesthetised with isoflurane, and sacrificed by cervical dislocation. Eyes were enucleated with a dorsal orientation cut and hemisected in carboxygenated (95% O₂, 5% CO₂) ACSF containing (in mM): 125 NaCl, 2.5 KCl, 2 CaCl₂, 1 MgCl₂, 1.25 NaH₂PO₄, 26 NaHCO₃, 20 glucose, and 0.5 L-glutamine (pH 7.4), supplemented with 0.1 μM Sulforhodamine-101 (SR101; Invitrogen) to identify blood vessels and damaged GCL cells. ACSF was continuously perfused at 4 ml/min and maintained at ∼36°C. Retinae were bulk-electroporated with the calcium indicator Oregon-Green 488 BAPTA-1 (OGB-1, 5 mM; Life Technologies) to label GCL neurons. The retina was flat-mounted ganglion cell layer (GCL)-side up onto an Anodisc™ (0.1 μm pore, 13 mm; Cytiva) and placed between two platinum disk electrodes (CUY700P4E/L, Nepagene/Xceltis). Nine pulses (∼9.2 V, 100 ms, 1 Hz) were delivered via a pulse generator/amplifier (TGP110/WA301, Thurlby Thandar/Farnell). The retina was then transferred to the recording chamber with the dorsal edge oriented away from the experimenter, allowed to recover for ∼30 min, and adapted to light stimulation using a binary dense noise stimulus (20×15 matrix, 40×40 μm² pixels, 5 Hz) prior to recording.

### Light Stimulation

#### Zebrafish imaging

Full-field light stimuli were delivered via a spectrally broad liquid light guide coupled to two CHROLIS 6-wavelength high-power LED light sources (Thorlabs). Across the two units, six LEDs were used, with peak wavelengths at 368 nm, 416 nm, 473 nm, 569 nm, 595 nm, and 627 nm. LED activation was synchronised to the scan retrace at a line rate of 500 Hz using ‘LED Zappelin’^123^. The output intensity of each LED was initially calibrated to 30 nW, which served as the maximum light level. For white stimuli, all six LEDs were combined, and their individual intensities were adjusted to approximate the spectral distribution of natural daylight in the zebrafish’s native habitat. The resulting intensities used in experiments were 3 nW (368 nm), 8.1 nW (416 nm), 14.1 nW (473 nm), 24 nW (569 nm), 26.7 nW (595 nm), and 30 nW (627 nm).

For retinal recordings, and the light guide was positioned adjacent to the animal at approximately 45° to ensure uniform illumination of the eye. For recordings of the arborisation field (AF) and optic tectum, larvae were mounted dorsally. In AF recordings, a single eye was presented with stimuli via a custom-built diffusion screen (63 gsm tracing paper, 3 cm × 1.9 cm, positioned 1.6 cm from the animal); for tectal recordings, both eyes faced the screen simultaneously. In both configurations, the light guide was positioned on the opposite side of the screen, ensuring that full-field illumination reached the animal exclusively through the diffusing surface.

For all experimental conditions, a constant background illumination period of at least 5 seconds was maintained following the onset of laser scanning prior to stimulus delivery. Full-field chromatic stimuli were presented as ‘white’ stimuli, in which all 6 LEDs were modulated on and off simultaneously. ‘White Chirp’ stimuli was presented as described before^1^.

#### Zebrafish Multielectrode array recordings

Eight spectral LEDs (λmax: 365 nm, 413 nm, 460 nm, 510 nm, 535 nm, 560 nm, 610 nm, 660 nm) were mounted onto an LED board (Adafruit 24-Channel 12-bit PWM) and driven by an LED driver (TLC5947, SPI Interface). LEDs 413-660 nm were bandpass filtered using Edmund Optics’ 34-493, 65-088, 65-095, 97-887, 65-103, 86-086, respectively. A liquid light guide (LLG5-8H, ThorLabs) was used to deliver light to a custom modified light projection engine (DLP LightCrafter E4500 MKII Fiber Couple) which projected light onto the RGC side of the retina.

All LEDs were set to 45 nW, corresponding to photopic conditions. Stimuli were custom written in C++, uploaded via PlatformIO, and delivered using a microcontroller (Adafruit ESP32 Feather V2). The area of all stimuli used in this study was greater than that of MEA electrode array. The stimuli comprised of flashes of light with all eight LEDs driven together (“white”), at ten different contrast levels (100-10% in 10% steps, 2 s On, 2 s Off). Neutral density (ND) filters (absorptive ND filters of optical density: 1.0 (NE2R10B, NG4), 2.0 (NE2R20B, NG9), 3.0 (NE2R30B, NG9), ThorLabs) were placed in the light path. ND0 corresponds to no ND filter being present. The same stimuli were played at each ND filter level, progressing from ND3 to ND0.

#### Mouse

Visual stimuli were delivered via a DLP projector (LCr, DPME4500UVBGMKII, EKB Technologies) through the objective onto the photoreceptor layer. The projector was equipped with external band-pass filtered green and UV LEDs (green: 576 BP 10; UV: 387 BP 11; AHF/Chroma) and a dual-band filter (390/576, F59-003, AHF/Chroma) to optimise spectral separation of mouse M- and S-opsins. Both LEDs were synchronised with the microscope scan retrace. Stimulus intensity was calibrated to range from ∼0.5 (black) to ∼20 × 10³ P*s⁻¹ per cone for both opsins, with an additional steady ∼10⁴ P*s⁻¹ per cone present during scanning due to two-photon excitation of photopigments.

Two stimuli were used for GCL Ca²⁺ imaging: (1) a full-field chirp stimulus (700 μm diameter), (2) a bright moving bar (0.3×1 mm, 1 mm/s) in eight directions to assess direction and orientation selectivity. A baseline period of ≥30 s was recorded after scan onset before each stimulus to avoid laser-induced effects on retinal activity.

### 2-photon imaging

2-photon imaging during visual stimulation was performed with a Movable Objective Microscope (MOM)-type 2P microscope (Sutter Instruments/Science Products) equipped with a mode-locked Ti:Sa laser (Chameleon Vision-S, Coherent) tuned to 920 nm for GCaMP excitation (larval zebrafish) / 927nm for OGB1 excitation (mouse) and a water immersion objective (larval zebrafish: W Plan-Apochromat 924 20x/1.0 DIC M27, Zeiss; mouse retina: CF175 LWD×16/0.8W, DIC N2, Nikon). For image acquisition, we used the custom-written software ScanM running under IGOR Pro 6.3 for Windows (WaveMetrics). Recordings were taken at 128 x 64 pixels (2ms scan, larval zebrafish retina), 256 x 128 pixels (2ms scan, larval zebrafish AF), 256 x 256 pixels (2ms scan, larval zebrafish optic tectum), and 64 x 64 pixels (2ms scan, for mouse retina).

### Pharmacological Manipulation

#### Zebrafish

For pharmacological manipulation, we injected ∼4 nL of a solution containing different drugs into the anterior chamber of the retina or forebrain depending on the experimental purpose. The following drugs were used: 1) for AC blockage in the retina and inhibitory circuits blockage in brain, a drug cocktail containing 3 different drugs were used, the estimated final concentration in the extracellular space was 5 µM Gabazine (Sigma-Aldrich) as antagonist of GABA_A_ receptors; 5 µM TPMPA (Sigma-Aldrich) as antagonist of GABA_C_ receptors; 5 µM strychnine (Sigma-Aldrich) as antagonist of glycine receptors. 2) For blocking GABA receptors only, we used solution containing only Gabazine and TPMPA with the same concentration. 3) For Group III mGluR receptor blockage, L-AP4 (2.5 µM, Sigma-Aldrich) were used. 4) For blocking glutamate transporters, TBOA (5 µM, Hello-Bio) were used. 5) For blocking Kainate receptors, ACET (2.5µM,Tocris) were used.

#### Mouse

All used drugs were added to the carboxygenated, perfused ACSF solution 15 min prior to the second recording of the GCL scan fields. For the drugs, the respective concentrations were used: 20 µM gabazine plus 75 µM TPMPA (GABA block), or 1 µM strychnine (glycine block). The ACSF solution with and without drug application was always kept at ∼36°C.

### Multielectrode array recordings

MEA recordings were performed on a MEA2100-System (Multichannel Systems, Germany). The MEA chip was 256MEA100/30iR-ITO-pr (Multichannel Systems, Germany), consisting of 256 electrodes of 30 µm diameter, arranged in a 16x16 grid, with 100 µm interelectrode spacing. The chamber above the array was 6 mm deep. Data acquisition was performed using Multichannel Experimenter software (Multichannel Systems, Germany), set to a sampling frequency of 20 kHz.

### Zebrafish Imaging Data Analysis

#### Preprocessing

Where necessary, images were xy-registered using the registration function provided in SARFIA^2^ running under IGOR Pro 9.0. ROIs were defined automatically using three approaches depending on data type: for BC data, regions of interests (ROIs) were placed automatically after local thresholding of the recording stack’s standard deviation (typically >80) projection using the full white step data, followed by filtering for size and shape using custom-written script under Igor Pro 9.0 (WaveMatrics), as described previously^1^; for AF data, ROIs were defined automatically based on local image correlation over time^3^; for brain data, using cell-lab^4^ for soma segmentation combined with a response-correlation projection, retaining only ROIs falling within a single soma-sized segment with response coherence ≥0.1^5^.

Fluorescence traces for each ROI were z-normalised, using the time interval 2–6 s recording onset. A stimulus time marker embedded in the recording data served to align the Ca^2+^ traces relative to the visual stimulus with a temporal precision of 1 ms. Responses were up sampled to 1 kHz and averaged over 5 trials. A response quality index (QI) was computed following Ref^4^, which for each ROI and all of its repeats quantifies the mean of the response variance divided by the variance of the response mean. Traces with QI higher than a threshold (0.5 for brain data, 0.35 for other recordings) were used for further analysis.

#### Response Polarity Classification

For the bipolar cell recordings, we used reconstruction approach described before^1^ using custom Python (v3.12) scripts. In brief, we reconstruct the average step response using four temporal components, this yield four corresponding weights. The response polarity is determined by the following rules: A ROI was considered as displaying an On-response if the sum of the light-transient and light-sustained weights exceeded 3-SD. A ROI was considered as displaying an Off-response if either the sum of the light-transient and light-sustained weights was more negative than 3-SD, or if the sum of the dark-transient and dark-sustained components exceeded 3-SD. If by these criteria a ROI display both On- and Off-responses, it was counted as On-Off. ROIs failing to elicit either On-or Off-responses were counted as non-responsive. For the GABAergic disinhibition experiment where drug effects prolonged the On plateau, an additional area-under-curve (AUC) criterion was applied to the half-step phase (AUC threshold = 1.5) to capture On-Off responses.

For AF and brain recordings, traces were smoothed (Savitzky-Golay filter, Scipy, window_length = 11) where necessary, and the derivative was computed over a defined response window (0.5 s); values exceeding a threshold (typically 1) were classified as responsive, assigning each ROI to Off, On, On-Off, or non-responsive groups.

### Zebrafish Multielectrode Array Data Analysis

Spike sorting was performed using the Spyking Circus^124^ with parameters listed in Supplemental Table T3. This yielded a total of *n* = 562 units across experiments (275 and 287 units in experiments 1 and 2, respectively), including responses to additional stimuli not analysed further here.

Subsequent analyses were restricted to responses obtained under the 90% contrast condition of the 100-10% contrast series. For each unit, On and Off response components were quantified as the mean spike rate within a 500 ms window following the respective light step (onset or offset), minus the mean spike rate during the remainder of the trial (baseline). Units were included for further analysis if either On or Off activation exceeded a minimum response criterion of 4 Hz (corresponding to 2 spikes in the 500 ms window); negative activation values were set to zero.

This procedure was applied independently for each neutral density (ND) condition, yielding *n* = 170 (ND0), 110 (ND1), 42 (ND2), and 0 (ND3) responsive units. For each included unit, an On–Off index was calculated (On-Off)/(On+Off), where values of −1 and +1 correspond to purely Off- and On-dominated responses, respectively, and intermediate values indicate mixed response profiles.

### Mouse Data Analysis

The data was preprocess as described previously^6^. The full-step phase was extracted from full-field chirp responses, and response polarity was determined using the algorithm (AF and brain data) described above. In short, traces were first filtered by a QI threshold of 0.35, then smoothed (Savitzky-Golay filter, Scipy, window_length = 11) prior to calculation. For each trace, a response window (0.5 s) was selected to determine the responsiveness for both on and off stimulus periods. Traces were then sorted into four groups (On, Off, On-Off, and non-responsive) by calculating the derivative of the response window for both on and off phases: values exceeding a threshold (typically 1) were classified as responsive.

### Polarity visualisations

To visualise response polarity in 2-photon scans (e.g. Figure 3A), we generated ‘On’ and ‘Off’ images from stimulus-averaged recordings (cf. Supplemental Movies M1–4). For each condition, we first averaged the movie across repetitions, then averaged frames corresponding to the first 2 s following light onset (‘On’) or offset (‘Off’) and subtracted the mean of the preceding 1 s baseline. These greyscale images were multiplied by the scan’s quality-index (QI) projection (see below), which highlights regions with consistent stimulus-locked activity while suppressing non-responsive areas and spontaneous events.

The resulting ‘On’ and ‘Off’ images were mapped to RGB space (green for On; magenta for Off) and jointly equalised to enhance responsive regions. ‘Merged’ panels show the combined RGB image. In some cases, we additionally computed an On-Off contrast, defined as (On − Off)/(On + Off) from the pre-RGB images. This yields pixel values between -1 (Off; magenta) and 1 (On; green), with 0 (white) indicating either balanced On-Off responses or absence of responses. Accordingly, this representation primarily highlights the presence or absence of polarity-specific signals.

QI projections were computed analogously to the ROI-based quality index (QI; see above), but on a per-pixel basis.

To visualise polarity distributions across populations (e.g. Fig. 2E), we randomly subsampled typically n = 750 ROIs per condition and arranged them as waterfall plots, sorted by IPL depth (retina) or response polarity (brain, mouse). Traces were colour-coded by assigned polarity: On (green), Off (magenta), On-Off (yellow), and non-responsive (light grey). Subsampling ensured comparable visualisation across experiments with different sample sizes. For Figure 6, smaller subsets were used (n = 500 in D; n = 250 in G) due to limited data.

### Transcriptomics Data Analysis

All transcriptomic datasets used in this study were downloaded from Ref^70^, which also includes datasets further associated with Refs^69,71,110,125^. Throughout we used bipolar cell clusters and their polarity-identities as defined in Ref^70^ and extracted their associated glutamate-receptor expression patterns. Because gene names, orthologues, and expression patterns are not perfectly aligned across species, the terms kainate, AMPA, mGluR, and EAAT refer here to the most relevant gene sets in each species (see Supplemental Table T1 for correspondence).

‘Expression contrast’ was computed as (𝐴 − 𝐵)/(𝐴 + 𝐵), where 𝐴 and 𝐵 represent the combined contribution of the respective receptor groups, defined as the product of mean scaled expression and fraction of cells expressing the transcript. For example, the Off-expression contrast (Figure 1C) was calculated with 𝐴 = AMPA_%expressed_ × AMPA_scaled expression_ and 𝐵 = Kainate_%expressed_ × Kainate_scaled_expression_. By this definition, values of −1 and +1 indicate exclusive dominance of kainate and AMPA receptor expression, respectively, whereas values near 0 reflect balanced contributions of both receptor types.

For Principal Component Analysis (PCA; Fig. 1G–L), we constructed a feature matrix in which rows correspond to bipolar cell (BC) clusters pooled across species (n = 246), and columns correspond to receptor features. For each of four receptor classes (kainate, AMPA, mGluR, EAAT), we included two measures: (i) mean scaled expression and (ii) fraction of cells expressing the gene (% expressed), yielding eight features in total. Because % expressed is inherently comparable across datasets, whereas scaled expression is not, we first normalised expression values within each species. Specifically, for each receptor feature, scaled expression values were z-normalised across BC clusters within species prior to pooling across species. This step mitigates species-specific differences in expression scaling. Following pooling, all features (both % expressed and scaled expression) were z-normalised across BC clusters to ensure equal weighting in the PCA. PCA was then performed on the resulting matrix using standard methods.

### Quantification and Statistical Analysis

Sample sizes were not predetermined statistically; randomization and blinding were not applied given the exploratory nature of the study. For comparisons between two groups, statistical comparisons were performed using Shapiro-Wilk tests to determine normality, then based on the results, we performed either paired/unpaired t-tests (normal distribution) or Wilcoxon rank-sum test (non-normal distribution, unpaired data) or Wilcoxon signed-rank tests (non-normal distribution, paired data). For comparisons across three or more groups, Kruskal-Wallis H-test followed by Mann-Whitney U test with Bonferroni correction were used (Fig. 8; OnOff-Index histogram comparisons). Details of the statistical comparisons for each figure are provided in Supplemental Table T2.

## Acknowledgements

We thank George Kafetzis and Marion Sillies for helpful discussions. **Funding** was provided by the Wellcome Trust (Investigator Award in Science 220277/Z20/Z to TB), the European Research Council (ERC-StG “NeuroVisEco” 677687 and ERC-AdG “Cones4Action” covered under the UK’s EPSRC guarantee scheme EP/Z533981/1 to TB), UKRI (BBSRC, BB/R014817/1 and BB/W013509/1 to TB), the Leverhulme Trust (PLP-2017-005, RPG-2021-026 and RPG-2-23-042 to TB), the Lister Institute for Preventive Medicine (fellowship to TB), The Swiss National Science Foundation (310030_204648 to SN), and the German Research Foundation (DFG; CRC 1233, project # 276693517; EU 42/12-1/RetNet4EC, project # 505379160 to TE). This research was funded in part by the Wellcome Trust (220277/Z20/Z). To promote Open Access, the authors have applied a CC BY public copyright licence to any Author Accepted Manuscript version arising from this submission.

## Author contributions

XW: Conceptualisation; Methodology; Investigation; Formal analysis; Data curation; Visualisation; Writing – original draft; LS: Investigation (adult zebrafish); Methodology; CF: Investigation (zebrafish brain neurons); Methodology; DG: Investigation (mouse retinal ganglion cells); Methodology; MS: Methodology (MEA hardware and analysis pipelines); TS: Resources (mouse protocols); SN: Resources (fish mutants); Funding acquisition; TE: Resources (mouse work); Funding acquisition; Supervision; TB: Conceptualisation; Supervision; Funding acquisition; Formal analysis; Visualisation; Writing – original draft. All authors: Writing – review & editing. The authors declare no conflict of interest.

---

**Supplemental Figure 1 Related to Figure 1.**
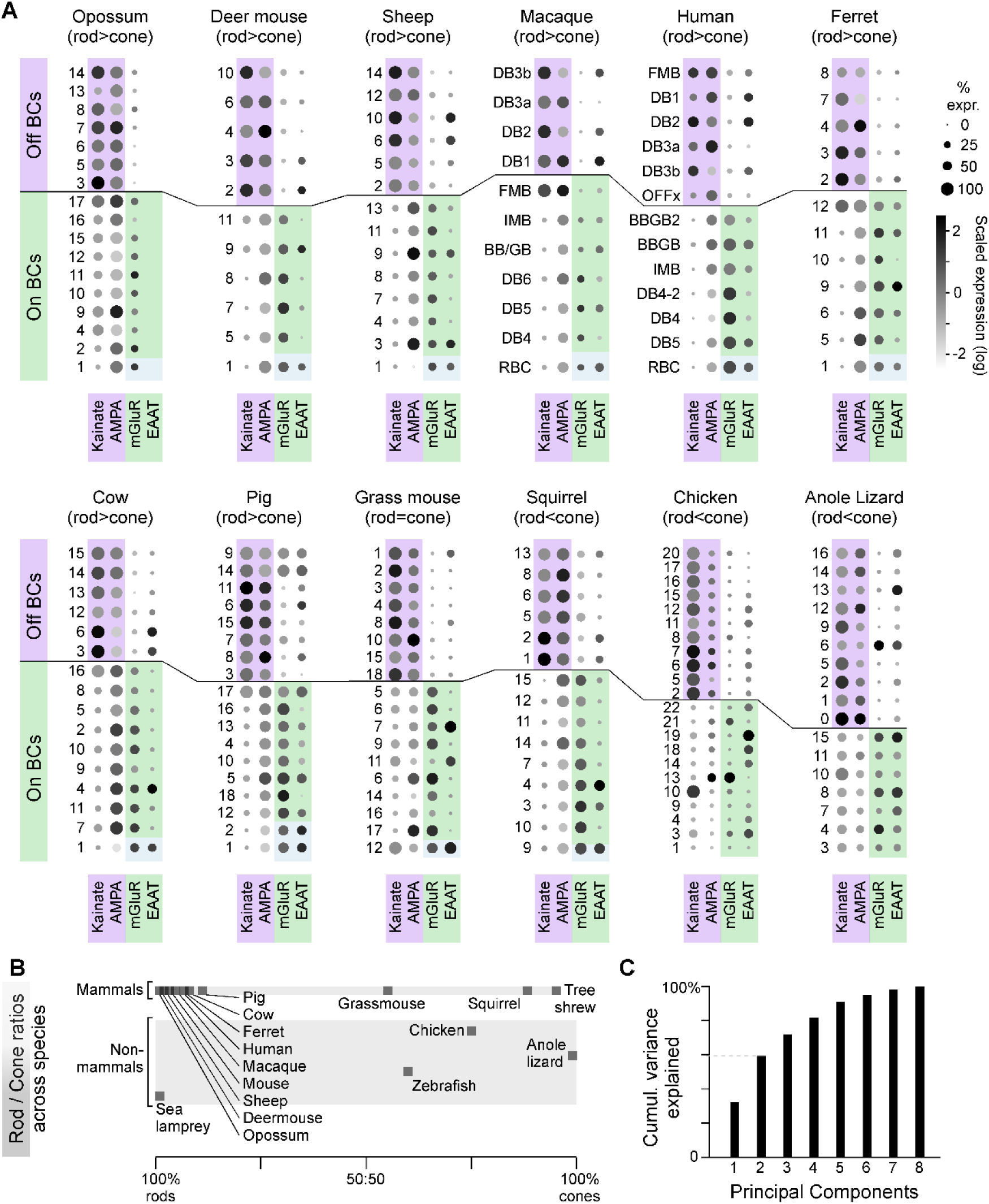
A, As in Figure 1B, summary of glutamate receptor expression patterns in BCs across species. **B,** Overview of typical rod:cone ratios across species used, based on the same transcriptomic datasets analysed in Figure 1. **C,** Cumulative variance explained across principal components (cf. Figure 1G–L).

**Supplemental Figure S2. Related to Figure 2.**
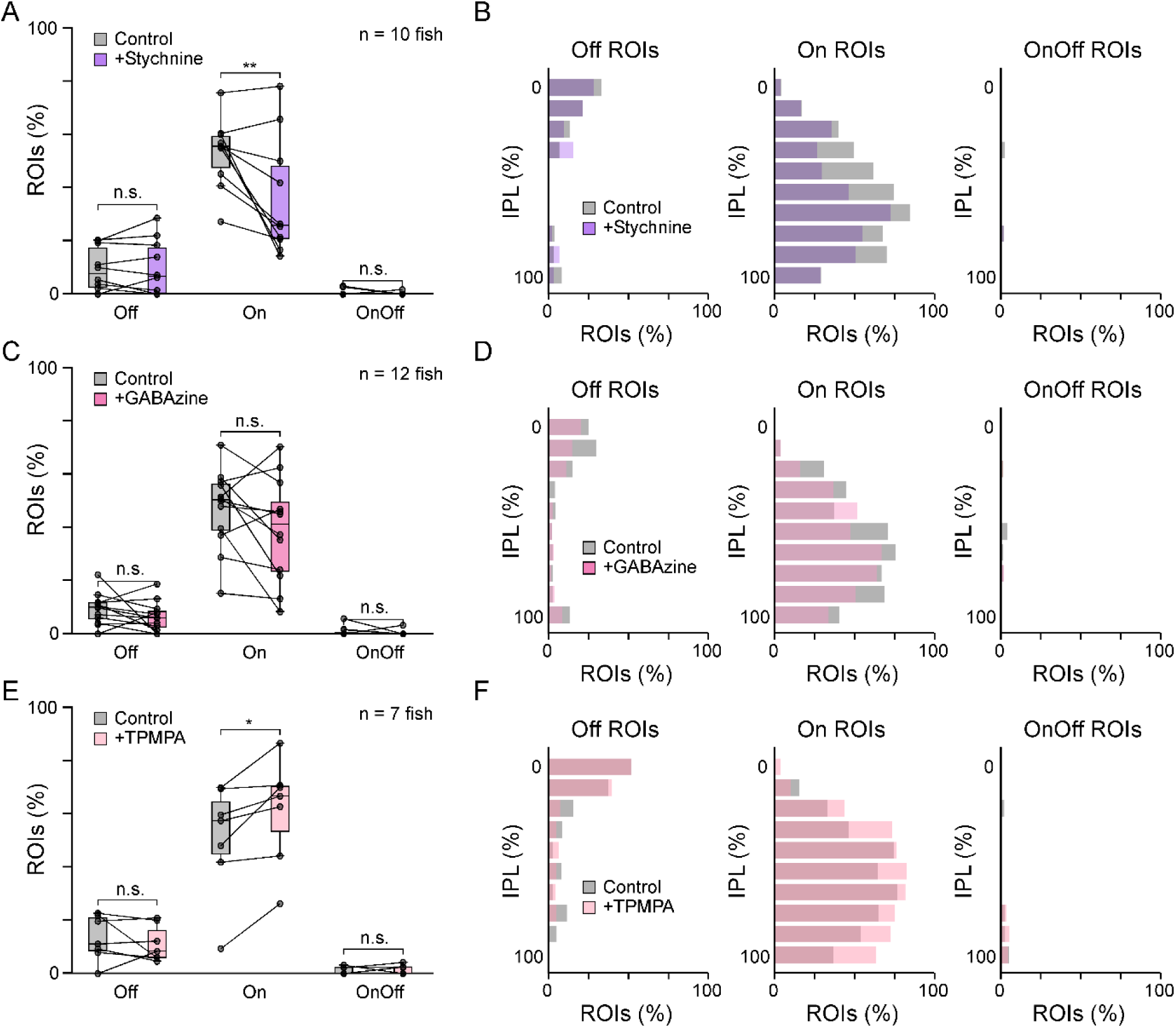
As in Figure 2F,G, for individual block of glycine receptors (**A,B**; n = 372 ROIs control and 399 ROIs drug, n = 10 fish), GABAA receptors (**C,D**; n = 601 ROIs control and 566 ROIs drug, n = 12 fish), and GABAC receptors (**E,F**; n = 375 ROIs control and 396 ROIs drug, n = 7 fish). These single-drug datasets were recorded in a different GCaMP line (ribeye-Gal4;UAS-syGCaMP3.5), which appears to show a weak expression bias toward On-stratifying BCs. Statistical comparisons use Wilcoxon signed-rank tests (see Supplemental Table T2).

**Supplemental Figure S3. Related to Figure 3.**
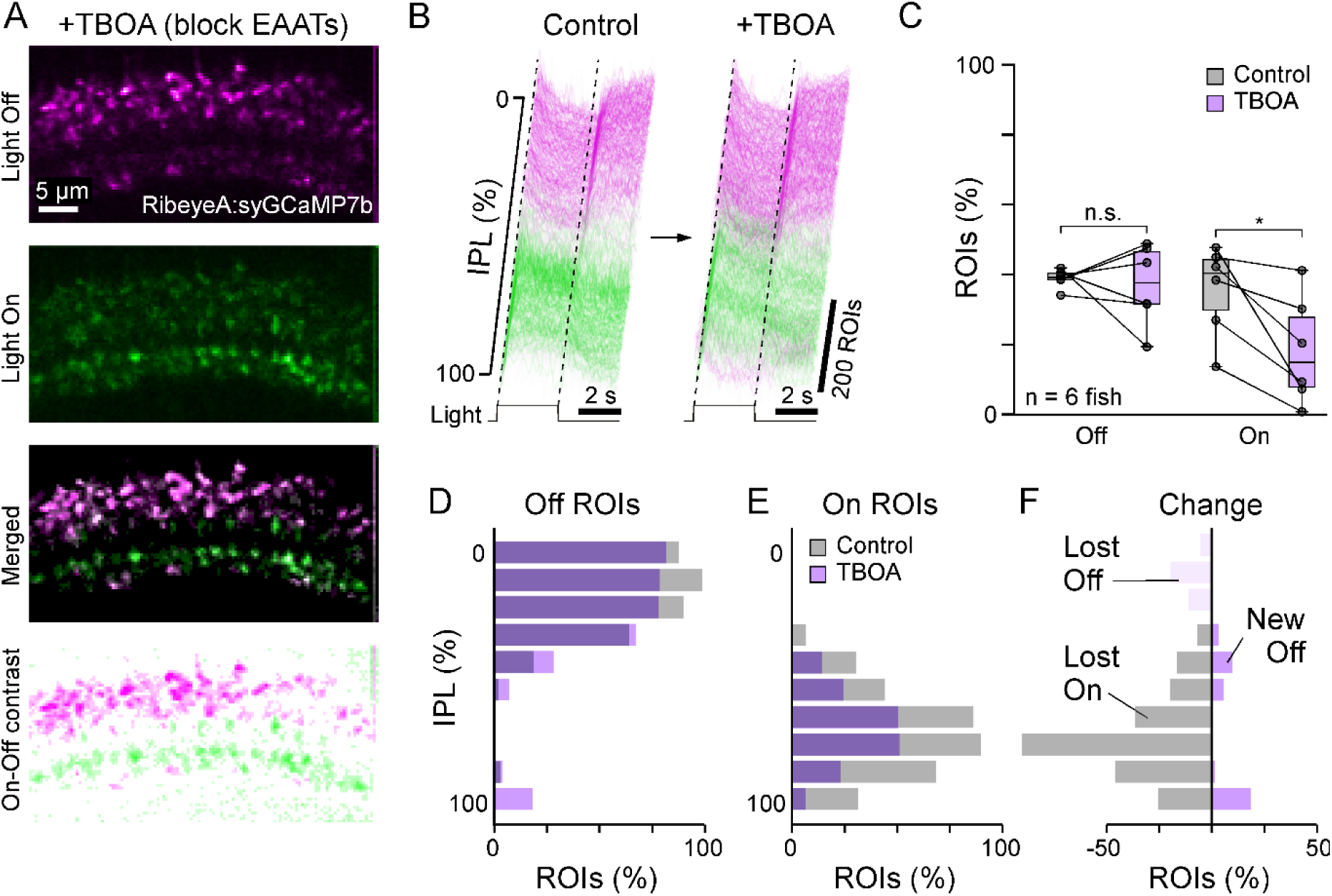
A–F, As in Figure 3F–J, here using TBOA to broadly block EAATs. Although weaker, the effects qualitatively mirror those observed in EAAT5b^-/-^ mutants.

**Supplemental Figure S4. Related to Figure 7.**
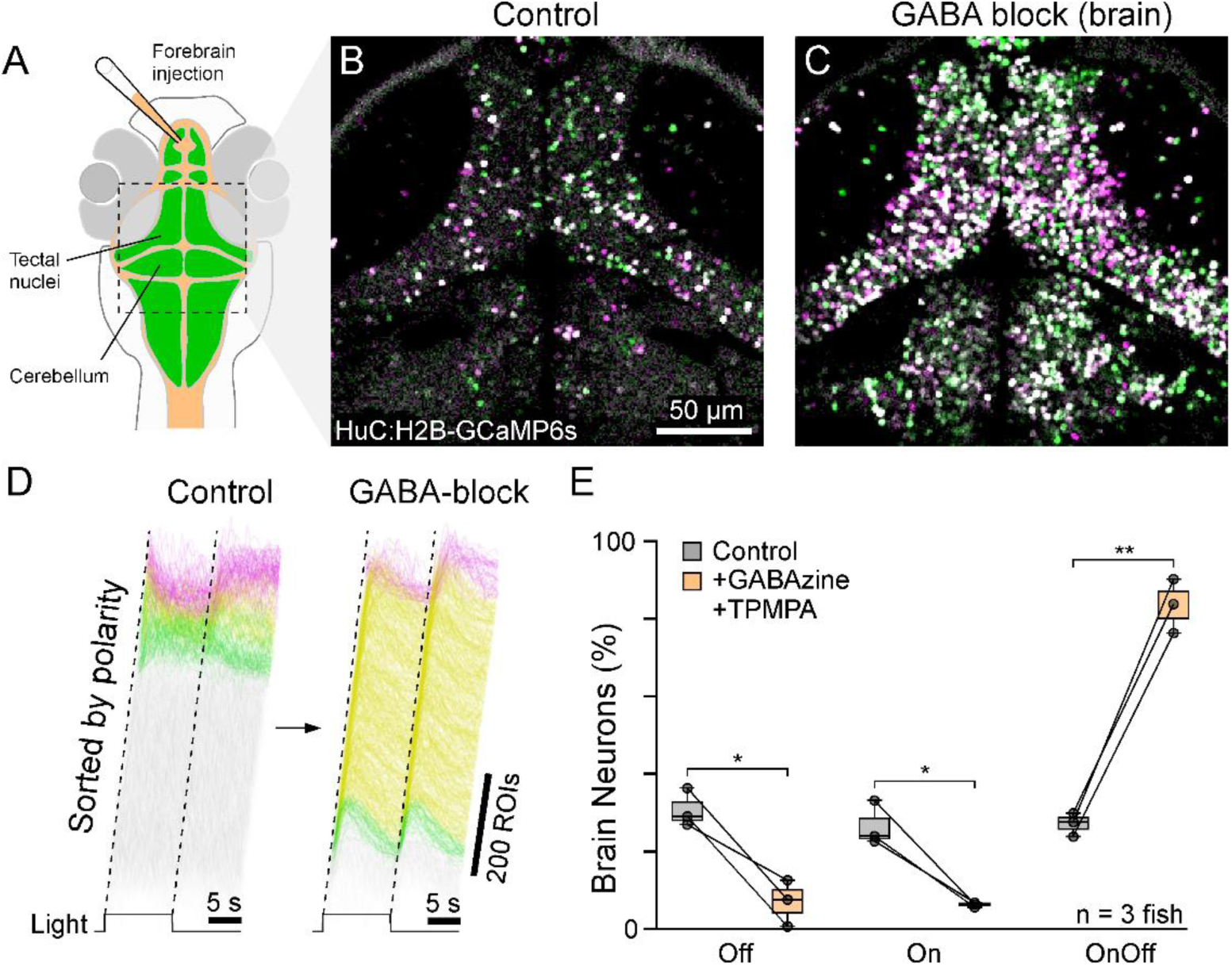
As in Figure 7F–J, but selectively disrupting GABAergic transmission while leaving glycinergic circuits intact. n = 3,405 ROIs “control” and n = 5,525 ROIs “drug”, from 3 fish. Statistical comparisons use paired t-tests (see Supplemental Table T2)

