## Supplementary figures and images for "On-Off coding is latent in vertebrate visual circuits"

### Supplemental Movie M1

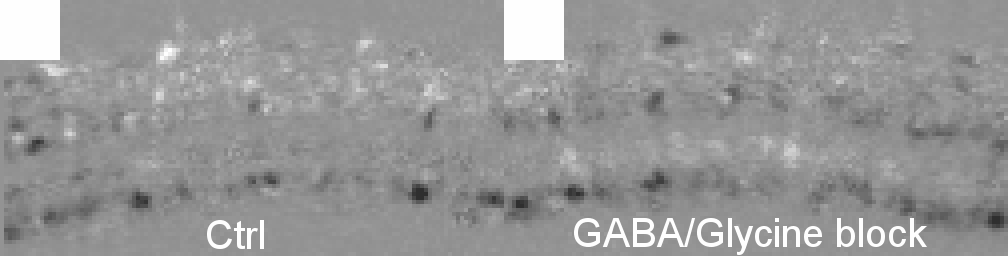

### Supplemental Movie M2

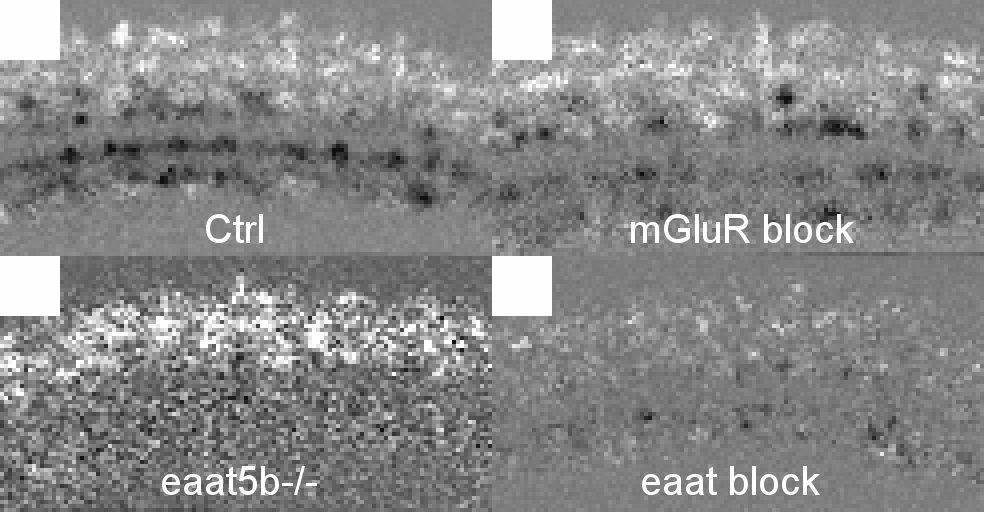

### Supplemental Movie M3

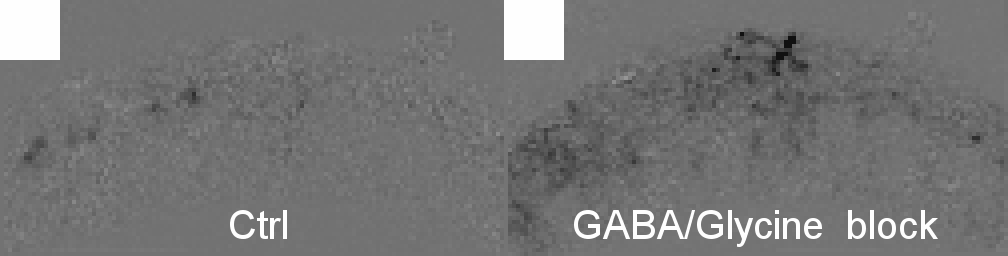

### Supplemental Movie M4

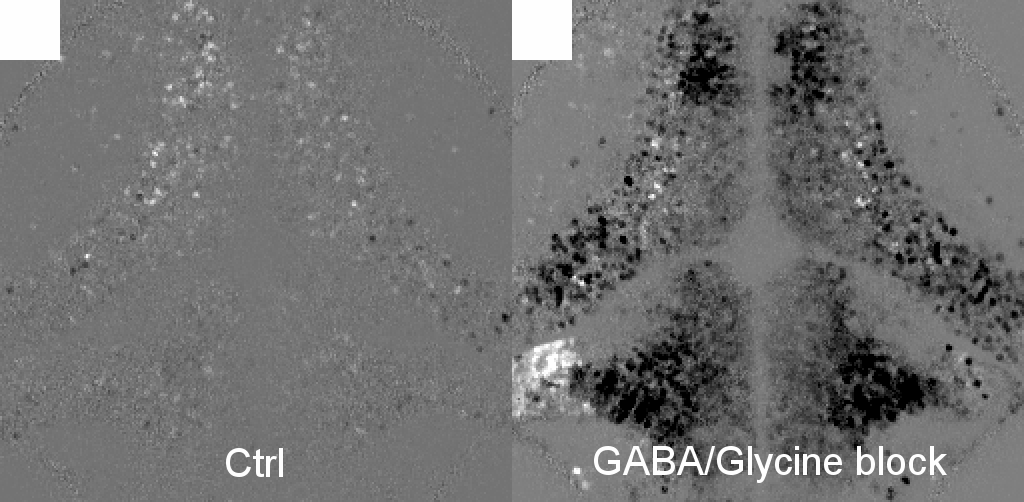
